# A multi-pore model of the blood-brain barrier tight junction strands recapitulates the permeability features of wild-type and mutant claudin-5

**DOI:** 10.1101/2025.01.29.635460

**Authors:** Alessandro Berselli, Giulio Alberini, Linda Cerioni, Fabio Benfenati, Luca Maragliano

## Abstract

In the blood-brain barrier (BBB), endothelial cells are joined together by multi-protein assemblies called tight junctions (TJs), which seal the paracellular space and restrict the passage of substances. Among the proteins forming BBB TJs, Claudin-5 (Cldn5) is the most abundant one. Structural models for complexes of Cldn5 and other claudins have been proposed and assessed via experimental and computational approaches. In these models, first introduced for channel-forming, selectively permeable claudins, protomers are arranged to form pores in the paracellular space that regulate transport by electrostatic and/or steric effects arising from the pore-lining residues. With limited exceptions, however, previous computational studies focused on oligomers of only a few subunits, while extended polymeric claudin strands form TJs. Here, we employ multi-microsecond all-atom molecular dynamics and free energy (FE) calculations to study two distinct models of TJ-forming Cldn5 complexes, called multi-Pore I and multi-Pore II, each comprising sixteen protomers arranged around three adjacent pores. Free energy calculations of water and ions permeation across the pores reveal that, in both models, the passage of ions is hindered by FE barriers, which are higher than in single-pore architectures. Moreover, only the multi-Pore I structural model recapitulates the effect of the G60R variant of Cldn5, making it permeable to anions. The results provide new insights into Cldn5 structure and function and validate a structural model of BBB TJs that may be useful for the study of barrier impairment in brain diseases and for developing new therapeutic approaches.

## Introduction

The blood-brain barrier (BBB) is a protective interface that separates the capillary blood flow from the brain parenchyma, preventing the access of harmful substances to the brain. It is composed of pericytes, astrocytes and endothelial cells, whose lateral membranes are tightly bound by protein complexes named tight junctions (TJs)^1–3^. TJ proteins form networks of transmembrane (TM) strands in each cell that associate with those of the neighboring cells to seal the paracellular space between them and regulate transport. Since TJ dysfunctions have been associated with various disorders, understanding their structural properties is essential for developing effective clinical applications^4–7^.

Claudin-5 (Cldn5) is the most abundant component of the BBB strands. Its assemblies form the TJ backbone and strictly limit paracellular transport. This makes Cldn5 a promising target for the development of drugs and strategies for direct brain delivery^8^. While Cldn5 subunits are known to form *cis*- and *trans*-interactions within each cell and between neighboring ones, respectively, the fine details of the TJ complexes remain substantially unknown due to the absence of experimentally determined structures. In the last decade, several structural models of claudins assemblies were proposed, and investigated also for Cldn5^9–11^. Interestingly, some of them display pore cavities in the paracellular space, oriented parallel to the lateral membranes, even when representing barrier-forming TJs such as those of the BBB. Two of these models have been investigated thoroughly as single pores made of four claudin monomers, usually referred to as Pore I and Pore II^6^. The former originates from the first model of TJ architectures, suggested for claudin-15 (Cldn15) in Ref. 12 (called the Suzuki or joined anti-parallel double-row-JDR-model), whereas the latter was first proposed in Ref. 9 for Cldn5. While the single pore tetrameric assembly is clearly not extended enough to seal the paracellular space, it represents the minimal structural unit suitable to study the channel or barrier properties of the corresponding TJ, making it a convenient system for computational investigations. Indeed, all-atom (AA) molecular dynamics (MD) simulations revealed that when wild-type (WT) Cldn5 is considered, both single Pore I and Pore II pose free energy (FE) barriers to the permeation of ions, consistent with the paracellular hindrance function^13,14^. While the barrier heights are moderate for some ions, simulations also showed that Pore I, and not Pore II, could reproduce the experimentally known effect of the hemiplegia-causing Cldn5 G60R mutation,^15^ making the BBB TJs permeable to anions^16^. The validity of the Pore I topology, both at the single-^17–21^ and multiple-pore levels, ^22–26^ was also assessed for other members of the claudin family, homologous to Cldn5 and expressed in the TJs of different tissues. The Single-Pore II was recently used in different works ^27–29^ to simulate BBB opening by shock waves^30^.

Considering extended, multimeric assemblies is of utmost importance to advance our knowledge of the TJ architectures, as they better represent the physiological environment. ^22–26^ Indeed, higher-order claudin complexes include inter-subunit interactions that are not present in the tetramers and might require specific conformations of the two extracellular loops (ECL1 and ECL2). However, detailed studies of Cldn5 multi-pore systems are still lacking.

Here, we employed standard AA-MD to study two extended versions of the pore models (named multi-Pore I and multi-Pore II), each made of sixteen Cldn5 monomers arranged to form three adjacent (i.e., parallel) pores. Then, we used the umbrella sampling (US) method to calculate the FE profiles of single Na^+^ and Cl*^−^*ions or water molecule permeation through the pores for both WT Cldn5 and the G60R variant. Results show that, in both models, the passage of ions is hindered by FE barriers, which are higher than in single-pore architectures. Moreover, only the multi-Pore I structure recapitulates the pathogenic effect of the G60R mutation in Cldn5 by becoming attractive for anions.

Our results provide insight into the structural features of multimeric Cldn5 complexes, representing segments of the BBB TJs. They confirm the validity of the Suzuki model^12^ for claudin-based paracellular structures, even in barrier-forming TJs, and provide useful information for the development of drug delivery strategies to modulate the BBB paracellular permeability by targeting the protein-protein contact interfaces^31^.

## Results

The Cldn5 multi-Pore I and multi-Pore II models are shown in Figures 1 and 2, respectively. Each one of them comprises sixteen protomers (**Figure S1**) assembled to form three adjacent paracellular cavities, named Pore 1, 2 and 3. Conversely, the single-pore models investigated in our previous works^14,16^ are indicated as single-Pore I and single-Pore II.

**Figure 1.**
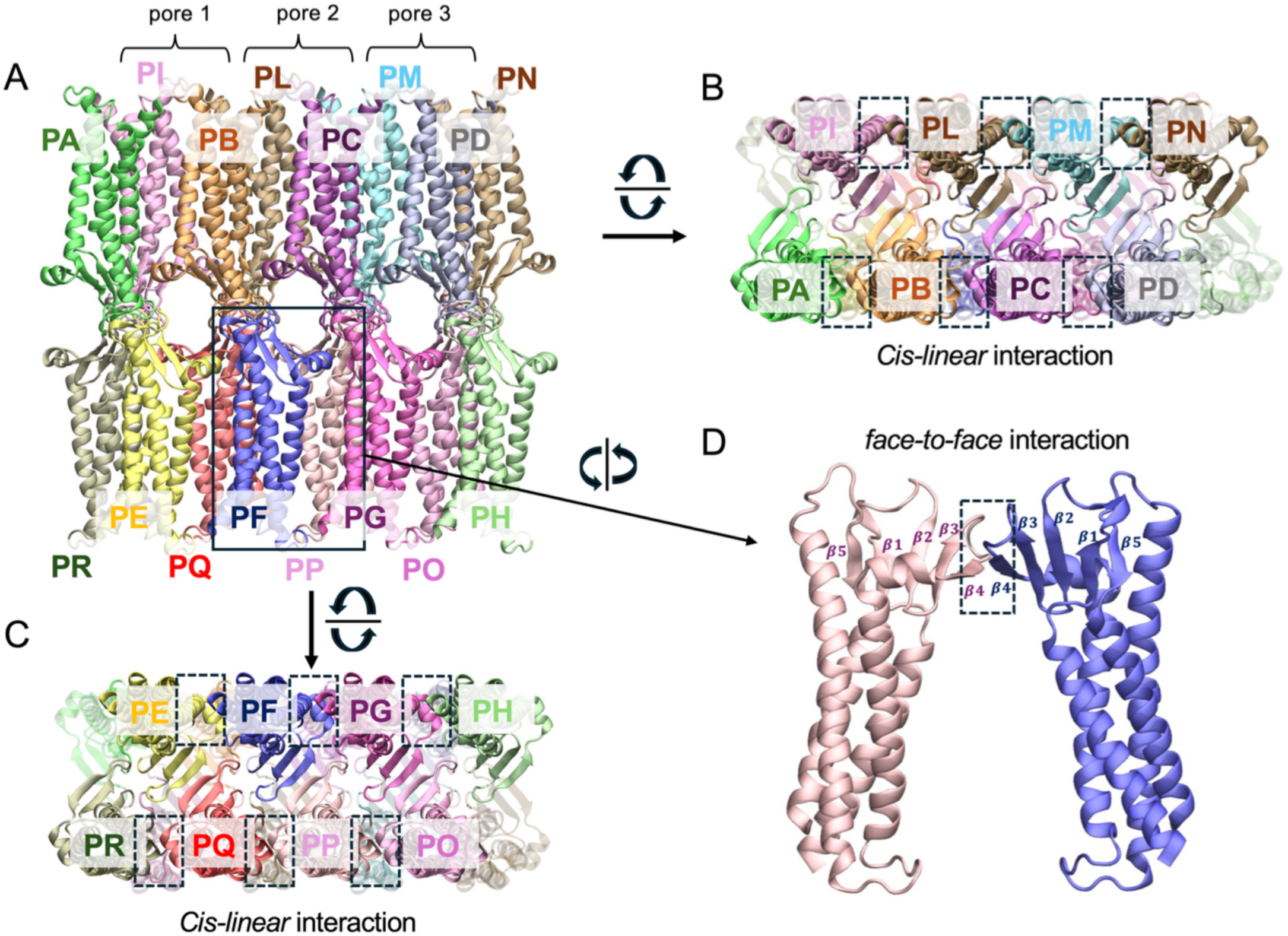
The architecture of the multi-Pore I model. The assembly includes sixteen Cldn5 monomers interacting via both cis- and trans-interfaces and forming three adjacent β barrel-like pores in the paracellular space. **A**, apical/basal view. **B**, **C**, lateral views, as seen from the cytosol of adjacent cells. Rectangles indicate protein domains involved in the cis-linear interface. **D**, two Cldn5 subunits interacting via the cis- face-to-face interface (highlighted by the rectangle).

**Figure 2.**
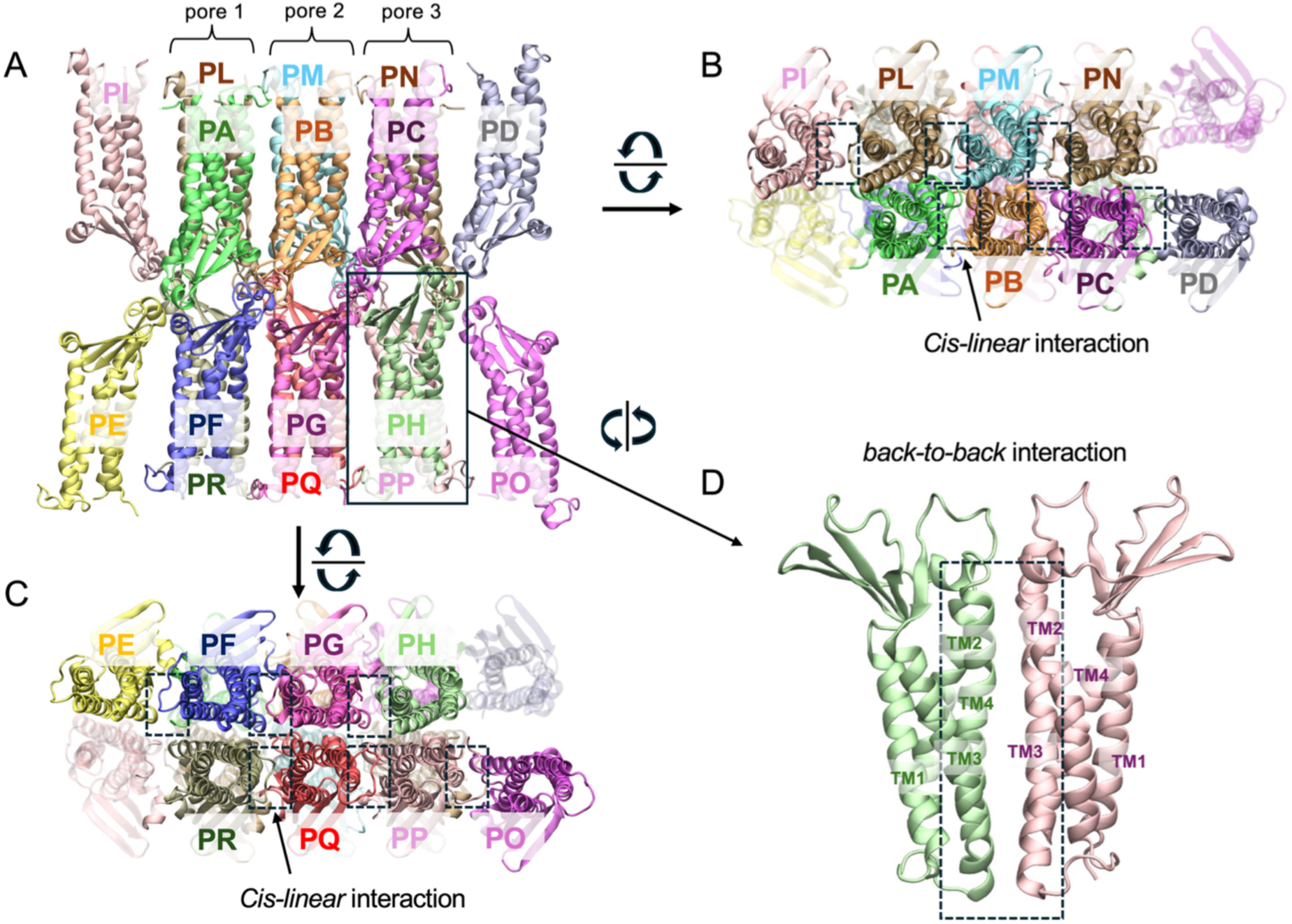
The architecture of the multi-Pore II model. The assembly includes sixteen Cldn5 monomers interacting via both cis- and trans-interfaces and forming three adjacent β-barrel-like pores in the paracellular space. **A**, apical/basal view. **B**, **C**, lateral views, as seen from the cytosol of adjacent cells. Rectangles indicate protein domains involved in the cis-linear interface. **D**, two Cldn5 subunits interacting via the cis- back-to-back interface (highlighted by the rectangle).

We simulated the multi-Pore I and II models using standard AA-MD and hydrogen mass repartition (HMR)^32–34^. We considered both WT and G60R Cldn5, and, for multi-Pore I, two versions of the variant structure differing by the orientation of the arginine side chains with respect to the pore lumen. These were named G60R-A and G60R-B, for outward- and inward-pointing chains, respectively. Positional restraints were applied to a few Cα atoms of the TM segments and of the ECLs of the peripheral protomers (extended set of restraints, see **Figure S2** and Methods section). We also ran two control simulations of the WT multi-Pore I using standard atomic masses and two distinct sets of positional restraints, the extended one and a restricted one comprising only Cα atoms of the peripheral protomers.

### Structural stability of the models from standard MD simulations

The time evolution of backbone Root Mean Squared Deviations (RMSDs), calculated for the MD simulations with the extended set of restraints, is reported in **Figure** 3. The atoms used for the calculation are illustrated in **Figure S3**. For the WT multi-Pore I (panel A), the RMSDs of the full system (ECLs plus TM domains, excluding peripheral protomers) and those of the individual pores (ECLs only) reached plateau values around 2 Å in the HMR trajectories (black lines), indicating stable conformations. Similar values were obtained in the control simulation with standard atomic masses (blue lines), revealing that the HMR setup does not affect results while saving computational resources. In the WT multi-Pore II (panel B), profiles showed a plateau at about 2 Å for the whole system and a stable central pore with an RMSD of ∼ 1 Å. Larger values were observed in the two lateral pores, but these more pronounced fluctuations were not associated with instabilities or large conformational changes in the architecture, as observed in more detail in the following. For the G60R multi-Pore I (panel C), the configuration with outward lateral chains (G60R-A) had RMSDs with a slow and limited drift. The RMSD of G60R-B (inward lateral chains) reached plateaus of about 1.75 A and 2.0 A in the whole paracellular domain and individual pores, respectively.

**Figure 3.**
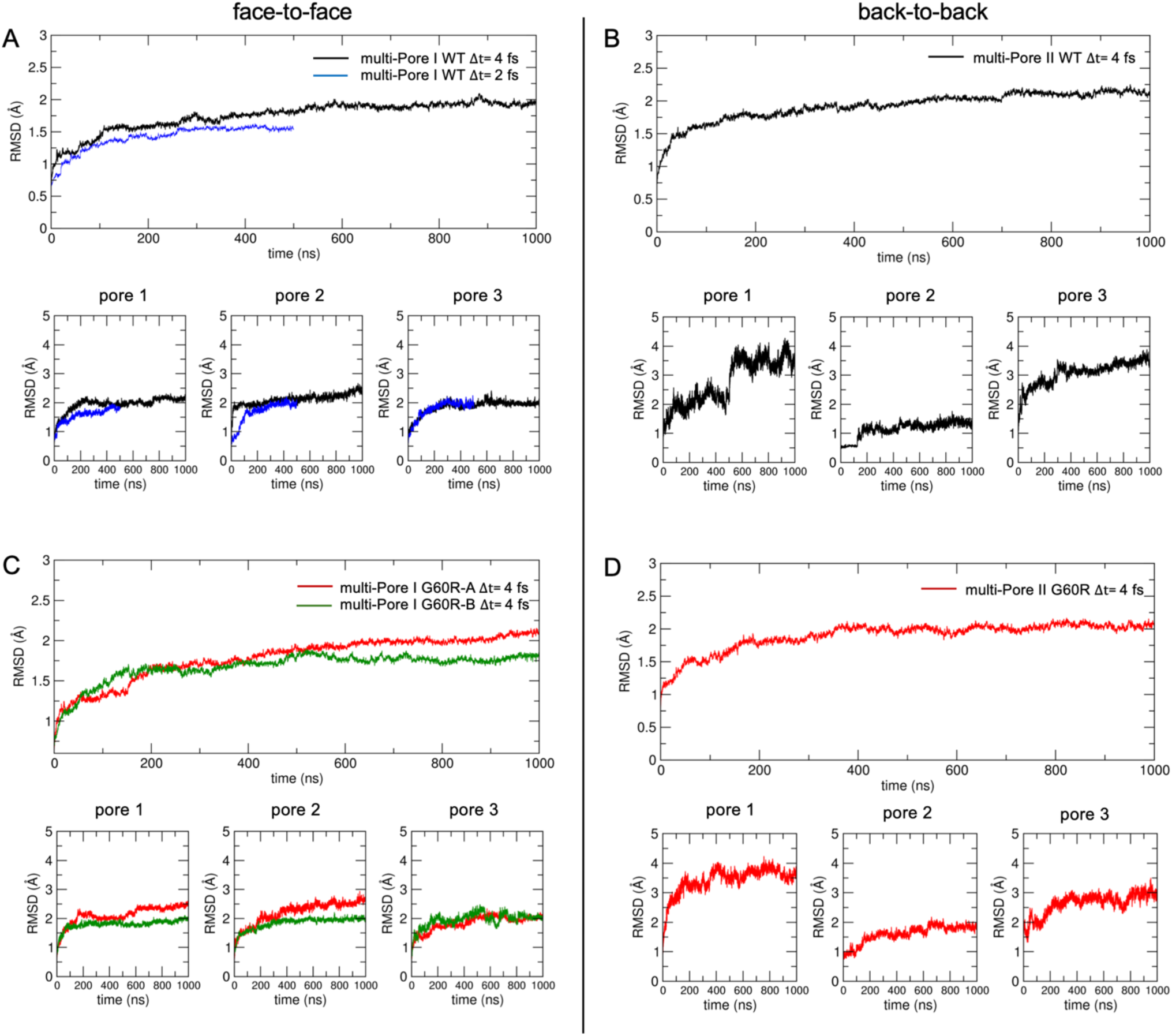
RMSD calculations. **A**, time evolution of backbone RMSD of multi-Pore I with HMR (black line) and standard masses (blue line), calculated for the whole system (ECLs plus TM domains) and the three separate pores (ECLs only, lower panels). **B**, backbone RMSD of multi-Pore II with HMR (black line) and standard masses (blue line), calculated for the whole system and the three separate pores (lower panels). **C,** backbone RMSD of multi-Pore I mutated models, whole system and individual pores (lower panels). **D,** backbone RMSD of mutated multi-Pore II, whole system and individual pores (lower panels).

Hence, in both conformations, RMSD values were comparable with the WT system, indicating that the mutation has a minor impact on the overall structure of the assembly. Finally, the RMSDs of the G60R multi-Pore II model (panel **D**) were stable at values close to the WT system, indicating a minor structural impact of the substitution.

#### Selection of a representative multi-Pore I model for the G60R variant

To proceed with the study of the G60R system using a unique multi-Pore I model, we compared the two versions by monitoring the conformations of the substituted arginine residues along the respective trajectories. We observed that over time, the arginine side chains gradually aligned in the two models, as revealed by the evolution over time of the distances calculated between their terminal carbon atoms (CZ) in diagonally opposed Cldn5 pairs (**Figure S4**). This result and those from the RMSD calculations above indicate that the two models are equivalent. Thus, we selected the G60R-A one for further analysis and refer to it as multi-Pore I G60R model.

#### Pore radius calculation

**Figure 4** shows pore radius profiles calculated over the AA-MD trajectories of multi-Pore I and multi-Pore II models in both WT and G60R configurations. The WT multi-Pore I model (HMR setup; panel **A**, black continuous line with gray area for standard deviation) had an average profile that is symmetric with respect to its center, where the four Q57 residues are located. It broadened at the peripheral openings, where it reached 6 Å. Along its axis, it displayed three constrictions of about 2.5 *−* 3 Å, one at the center and two symmetrically arranged, separated by two dilations of ∼ 4 Å. The control simulation with standard masses (blue dashed line) had a similar global profile, with the largest differences of ∼ 1 Å, i.e., of the order of standard deviations. When comparing these profiles to that of the single-Pore I that we calculated in Ref. 14 (black dotted line), we noticed that they have the same width at the center, although the latter features a more pronounced hourglass shape. In panel **B,** we report the average pore profile (solid line) of the multi-Pore II model (HMR setup). The structure had the same width at the center and the peripheries (radius ∼ 4.5 Å) and showed two constrictions of about 2.5 Å at *y ∼* 7.5 Å and 55 Å. In contrast, the single-Pore II structure (dotted line) was more uniform along the axis, with an average radius value of around 4 Å. In panel **C,** we show the pore radius of the multi-Pore I model built with the G60R variation (HMR setup). The profile reached ∼ 6 Å at the periphery and ∼ 2.5 Å at the center, like the WT system (panel **A**). However, the two additional restrictions observed in the WT structure were not present in the mutant. Finally, as for the WT, the mutated multi-Pore II structure had the same central width but a less pronounced hourglass shape than the corresponding single-Pore I^16^.

**Figure 4.**
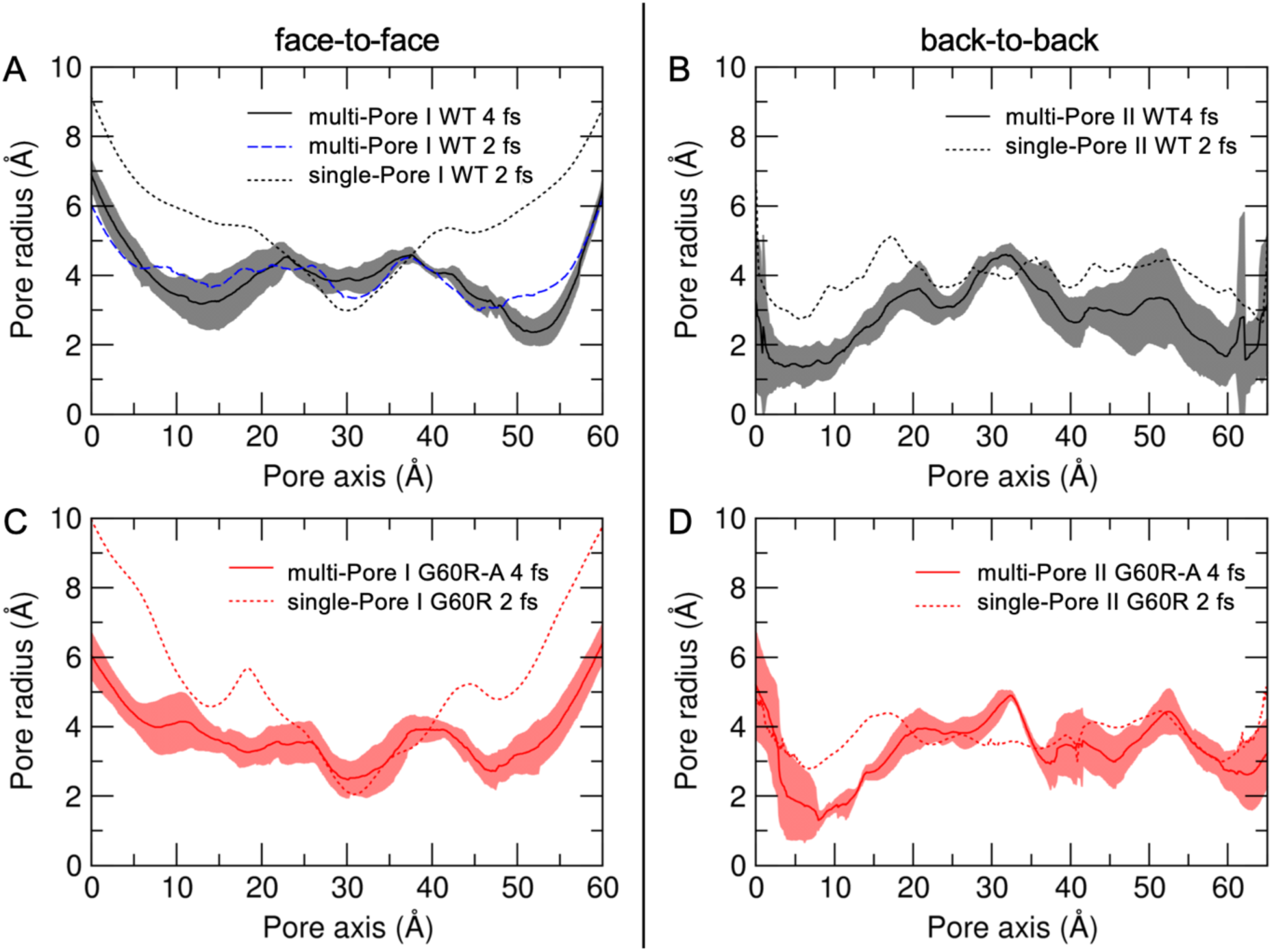
Pore radius profiles. **A**, WT multi-Pore I (black line), calculated as the average of the three pores from simulations with HMR or standard atomic masses (blue dashed line), and compared with the single-Pore I from Ref. 14 (black dotted line). **B**, WT multi-Pore II (black line), compared with the single-Pore II from Ref.14 (black dotted line). **C**, G60R multi-Pore I (red line), compared with the single-Pore I from Ref. 16 (red dotted line). **D**, G60R multi-Pore II (red line), compared with the single-Pore II from Ref. 16 (red dotted line).

In panel **D,** we report the pore radius of the G60R multi-Pore II (HMR), which is comparable to both WT (panel **B**) and the mutated single-Pore II from Ref. 16.

#### Stability of the linear interfaces

The high-order multimeric Cldn complexes here investigated include the so-called *cis-*linear interface (**Figure 5**), observed for the first time in the crystal lattice of Cldn15 and suggested to form key interactions stabilizing TJ strands^35^. To test the persistence of this interface in the simulations of our models, we monitored distances between residues in neighbor Cldn5 monomers, indicated in **Table 1**. We also monitored the distances between residues of *trans-*interacting monomers.

**Figure 5.**
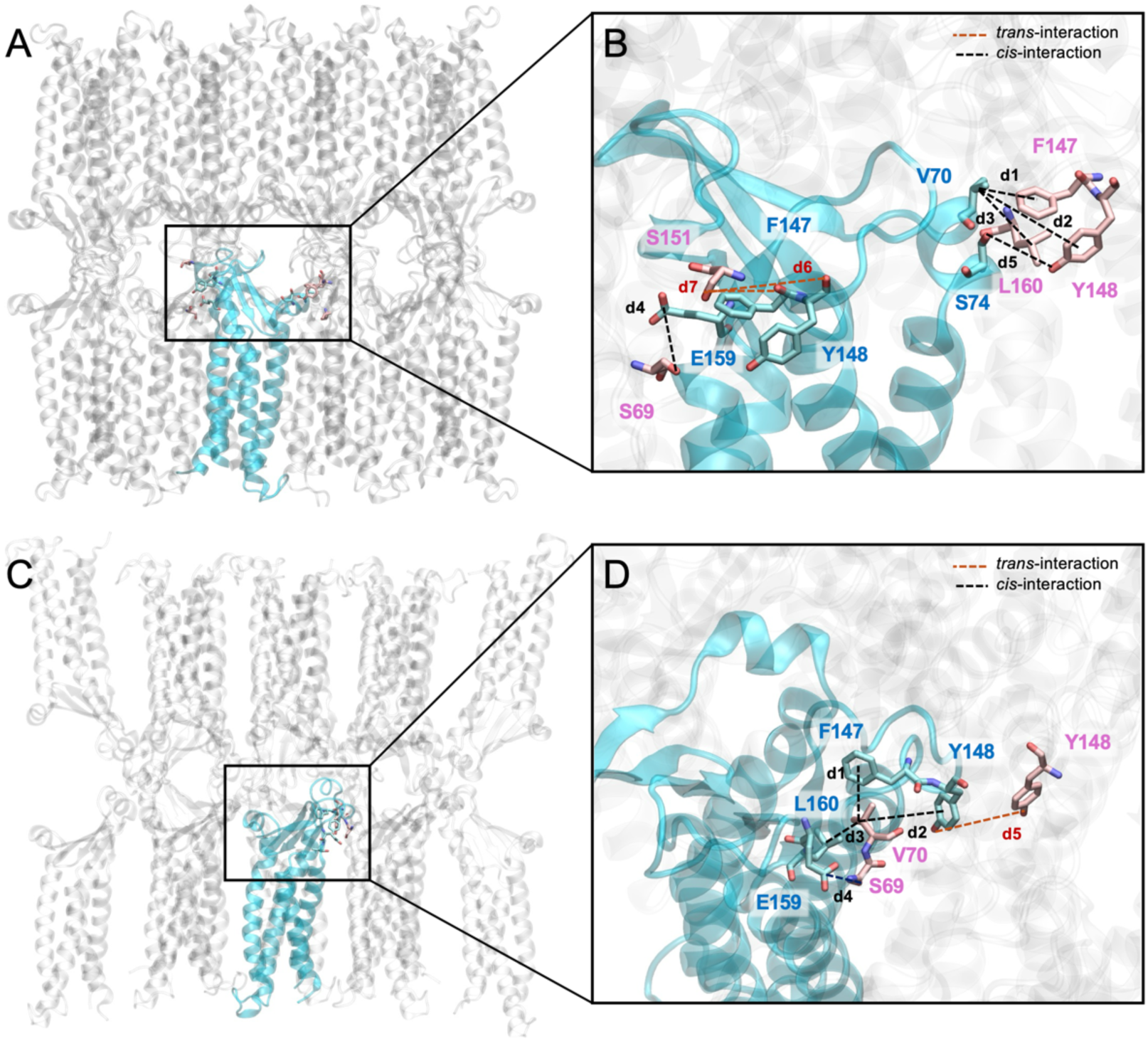
**A**, the network of the atomic interactions formed by a Cldn5 protein (in cyan) and the adjacent subunits in both the cis-linear and trans-interfaces of the multi-Pore I. **B**, magnification of the box in panel **A**. **C**, the network of the atomic interactions formed by a Cldn5 protein (in cyan) and the adjacent subunits in both the cis-linear and trans-interfaces of the multi-Pore II. **D**, magnification of the box in panel **C**.

**Table 1.** Distances defined to evaluate the stability of the cis-linear and trans-interactions. For each residue, we report the atom selected for the calculation, using CHARMM atom names (CB: C*β*, CD: Cδ, CG: Cγ, O: backbone oxygen-amide, OG: Oxygen γ-hydroxyl). COM indicates the center of mass of the carbon atoms forming the side chain of aromatic residues.

| Model | Cldn5 multi-Pore I | Cldn5 multi-Pore II |
| --- | --- | --- |
| d1 | V70 (CB) - F147 (COM) [*cis*-] | V70 (CB) - F147 (COM) [*cis*-] |
| d2 | V70 (CB) - Y148 (COM) [*cis*-] | V70 (CB) - Y148 (COM) [*cis*-] |
| d3 | V70 (CB) - L160 (CG) [*cis*-] | V70 (CB) - L160 (CG) [*cis*-] |
| d4 | E159 (CD) - S69 (OG) [*cis*-] | E159 (CD) - S69 (OG) [*cis*-] |
| d5 | S74 (OG) - Y148 (OG) [*cis*-] | Y148 (COM) - Y148 (COM) [*trans*-] |
| d6 | S151 (OG) - Y148 (O) [*trans*-] | - |
| d7 | S151 (OG) - F147 (O) [*trans*-] | - |
| d8 | S151 (OG) - S151 (OG) [*trans*-]<br>(control simulation only) | - |

In **Figure 6**, we report the values of these distances over time for the multi-Pore I model in both the WT (panel **A**, black curve) and the G60R (panel **B**) configurations (HMR setup). We also show values from the standard mass simulation (blue curve) of the WT model. Dashed green lines represent the values of the equivalent distances in the crystal lattice of Cldn15 (labeled as d1* to d4*) for reference. All distances were stable along the trajectories. In the WT, results for HMR matched those of standard masses, except for d4, for which the standard mass values were closer to the crystal one (green dashed line), with overlapping fluctuations in the two setups. Distances d1 to d3 were larger than their values in the crystal, with d2 showing the greatest difference, but this is not surprising since they all involve mobile side chain atoms. Inserting the mutation (panel **B**) did not affect distance values, except for d4, which better matches the crystal one. **Figure 7** reports the same analysis for the multi-Pore II model. Also in this case, all distances are stable along the trajectory, with the exception of d3, which, when the G60R mutation is introduced, switched from ∼7 Å to ∼ 12 Å.

**Figure 6.**
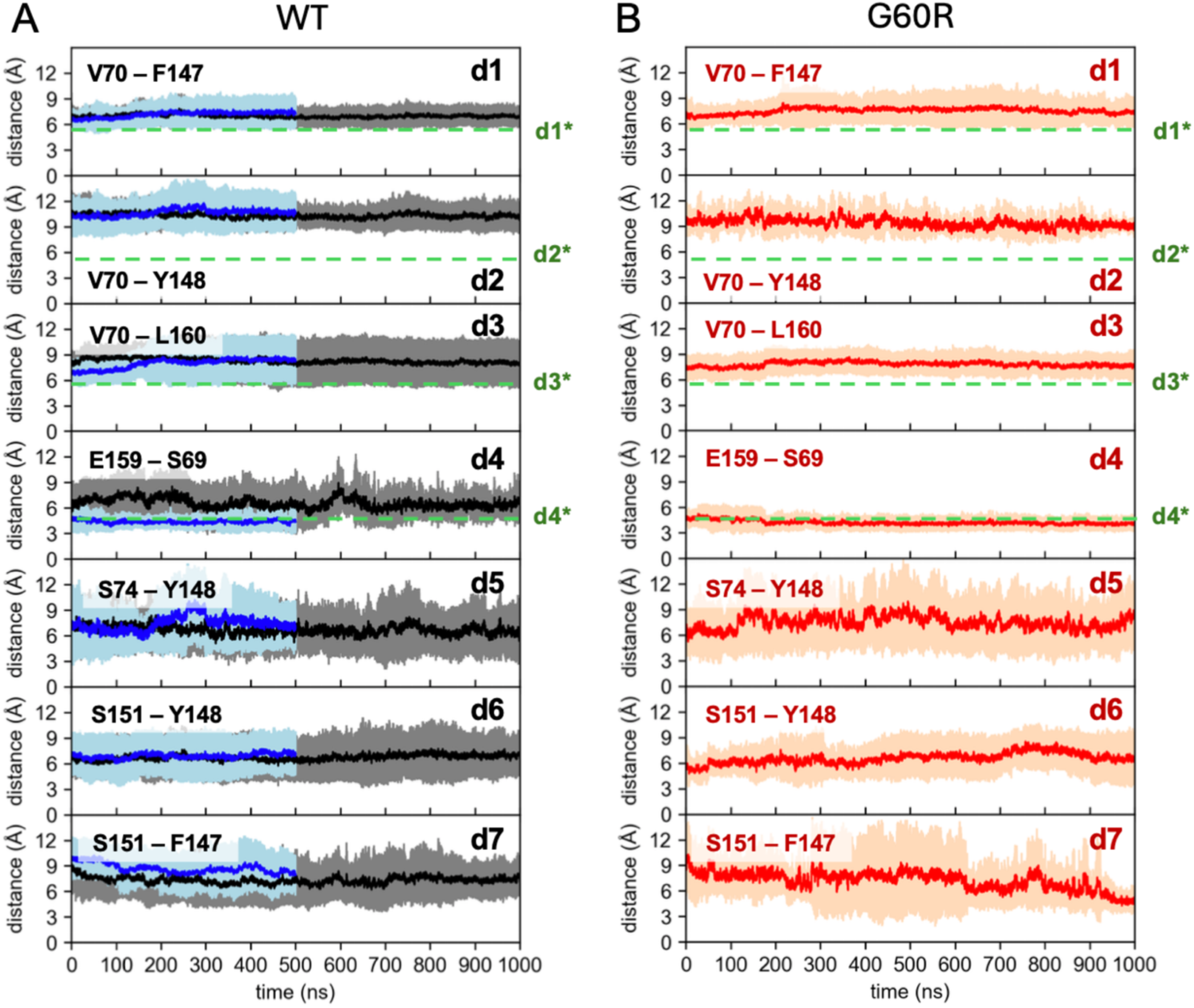
Time evolution of distances describing the cis-linear and trans-interactions for the WT (**A**) and G60R (**B**) multi-Pore I. Continuous lines and shaded areas are average values and standard deviations calculated over all realizations of the same distance in the assemblies using the HMR simulated trajectories. Blue lines are from the standard mass trajectories. Dashed green lines represent values observed in the Cldn15 crystal structure (PDB ID: 4P79).

**Figure 7.**
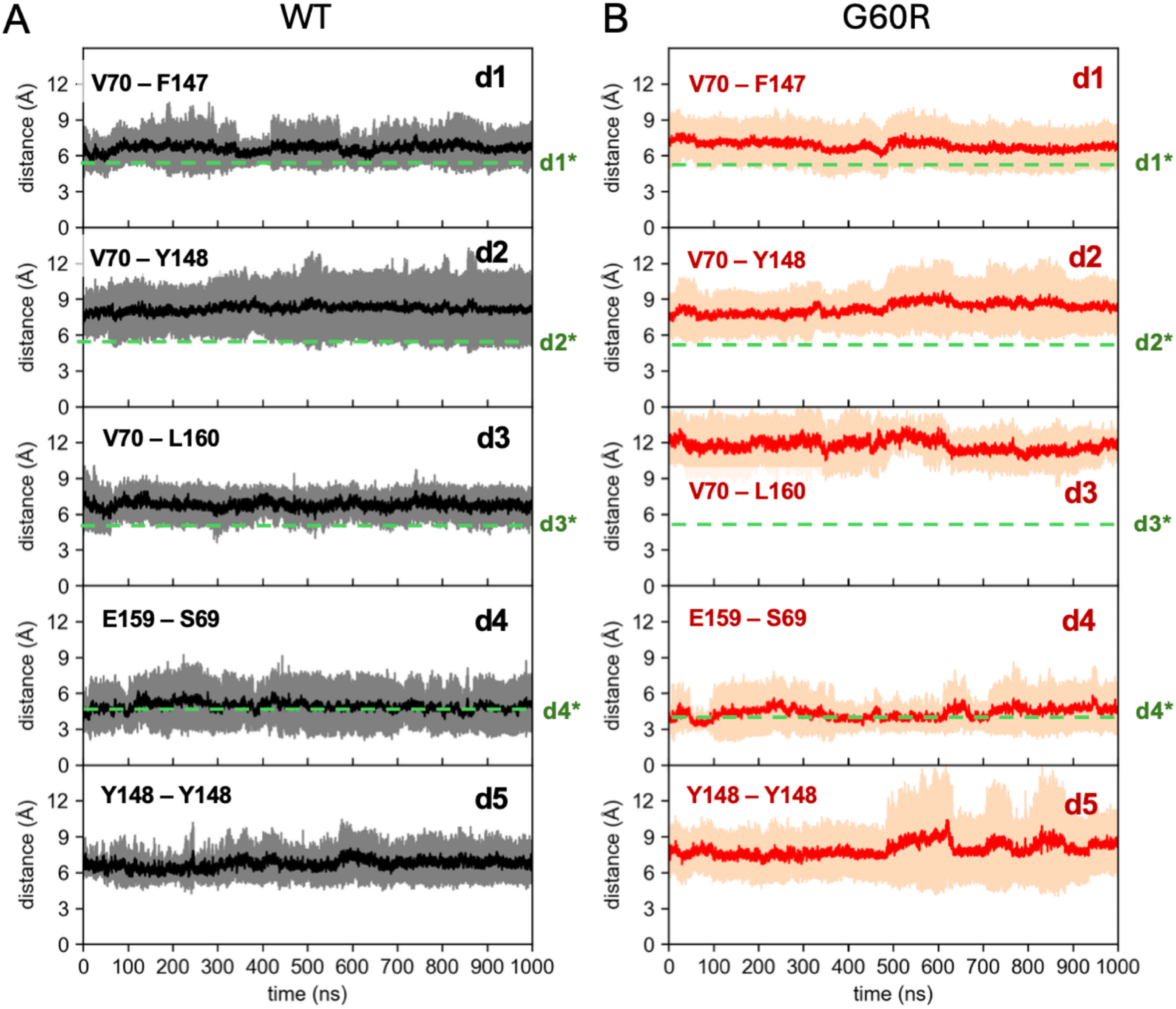
Time evolution of distances describing the cis-linear and trans-interactions for the WT (**A**) and G60R (**B**) multi-Pore II. Continuous lines and shaded areas are average values and standard deviations calculated over all realizations of the same distance in the assemblies using the HMR simulated trajectories. Dashed green lines represent values observed in the Cldn15 crystal structure (PDB ID: 4P79).

#### Control simulation of the WT multi-Pore I with a restricted set of restraints

To evaluate the effect of applying positional restraints to the complex, we performed a 500 ns-long control simulation of WT multi-Pore I with a restricted set of positionally restrained atoms, i.e., only some C*α* of the most peripheral monomers (**Figure S2A**, green spheres). Standard atomic masses were employed. In this trajectory, the backbone RMSD profile of the structure grew up to about 6.5 Å (**Figure S5**), and the drift was mostly due to a rearrangement of the TM helices since the RMSD of the ECL domains plateaued at around 4 Å. The same result was obtained for each individual pore. However, this reordering of the TM segments did not impact the *β* barrel structure of the pores, as can be observed by superimposing the multi-Pore I structure from the extended and restricted set of restraint trajectories at their respective 500 ns frames (**Figure S6**). More in detail, all the inter-monomer distances (d1 to d7) described above remained stable also in this system with fewer restraints. When comparing with the extended restraints set up (standard masses), only d6 and d7 showed values that differed, although with overlapping fluctuations (**Figure S7**). The additional distance d8, measured between S151 residues, differs more than the others between the extended and restricted set of restraint systems (**Figure S8**). However, by comparing configurations along the trajectories, we noted that the rearrangement of the residues involved in this distance does not alter dramatically the overall architecture of the complex. In the control simulation, the pore radius profile was again wider at the peripheries and narrower at the center, where it reached about 3.5 Å, but it was more uniform than for the other systems (**Figure S9**). To inspect the preservation of the pore topology in more detail, we monitored the time evolution of a set of cross-distances between residues from facing monomers, both at the periphery and in the middle of the cavity. The stability of these distances and their similar values in the extended and restricted restraints simulations (**Figure S10**) further confirm that the global structure of the complex is well preserved between the two systems.

#### Free Energy calculations of ion and water permeation across Cldn5 BBB strands

We computed 1D Free Energy (FE) profiles for the permeation of single physiological ions (Na^+^, Cl*^−^*) or a single H_2_O molecule through the central pore of the two structures (see **Figure S11** for an illustration of the pore and lining residues). Results for the multi-Pore I (both WT and G60R) are shown in **Figure 8**, while those of the multi-Pore II are in **Figure 9**, together with those previously obtained for single-pore architectures^14,16^. The flat profiles for water permeation indicate that both models are permeable to water, with the behavior of either WT or G60R Cldn5 that is similar to single-pore architectures.

**Figure 8.**
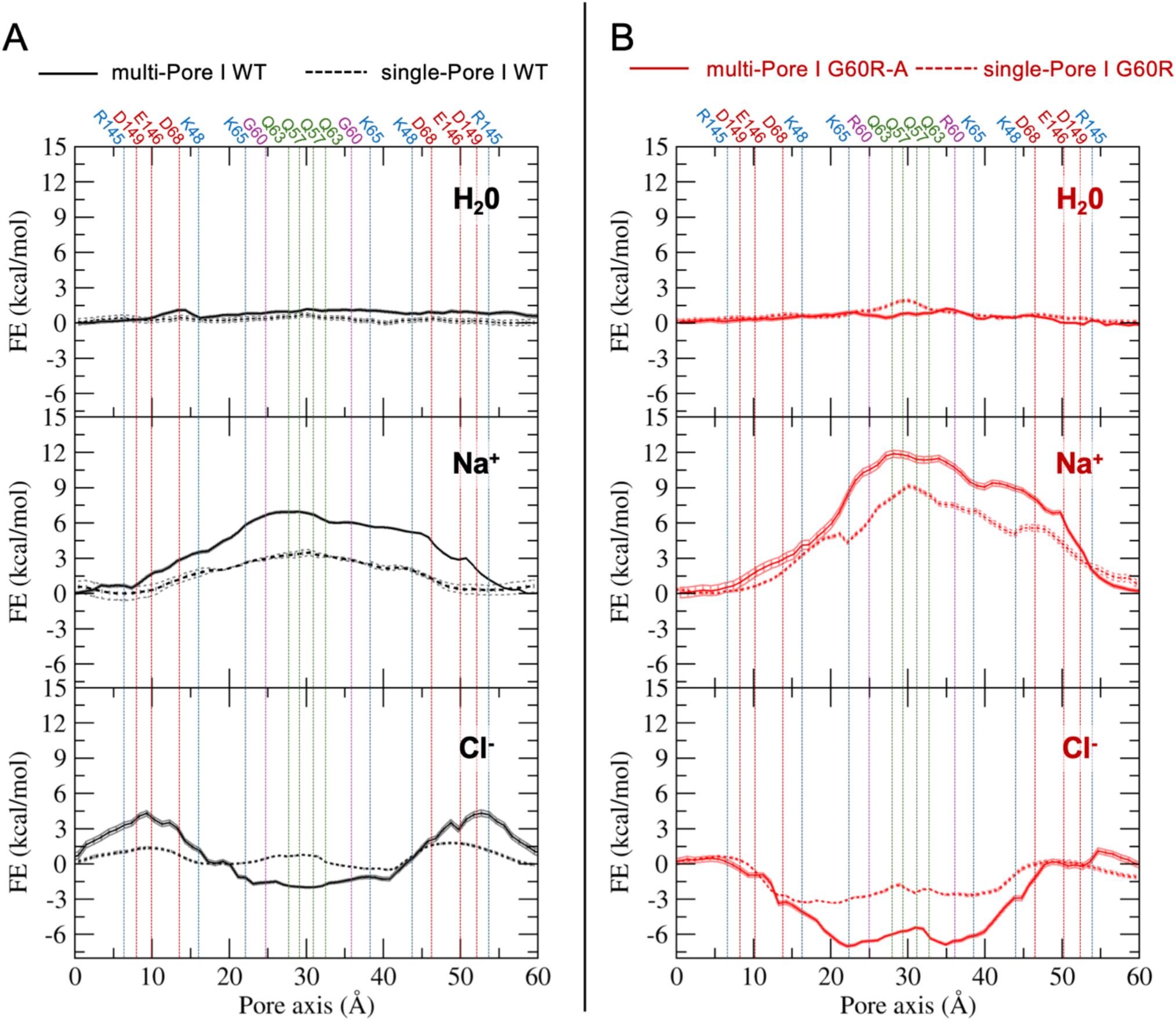
FE profiles for water and ion permeation across the BBB TJ models. **A**, WT multi-Pore I (solid black line), compared with single-Pore I^14^ (dashed black line). **B**, G60R multi-Pore I (solid red line), compared with single-Pore I^16^ (dashed red line). Vertical lines indicate the positions of the Cα atoms of the pore-forming residues.

**Figure 9.**
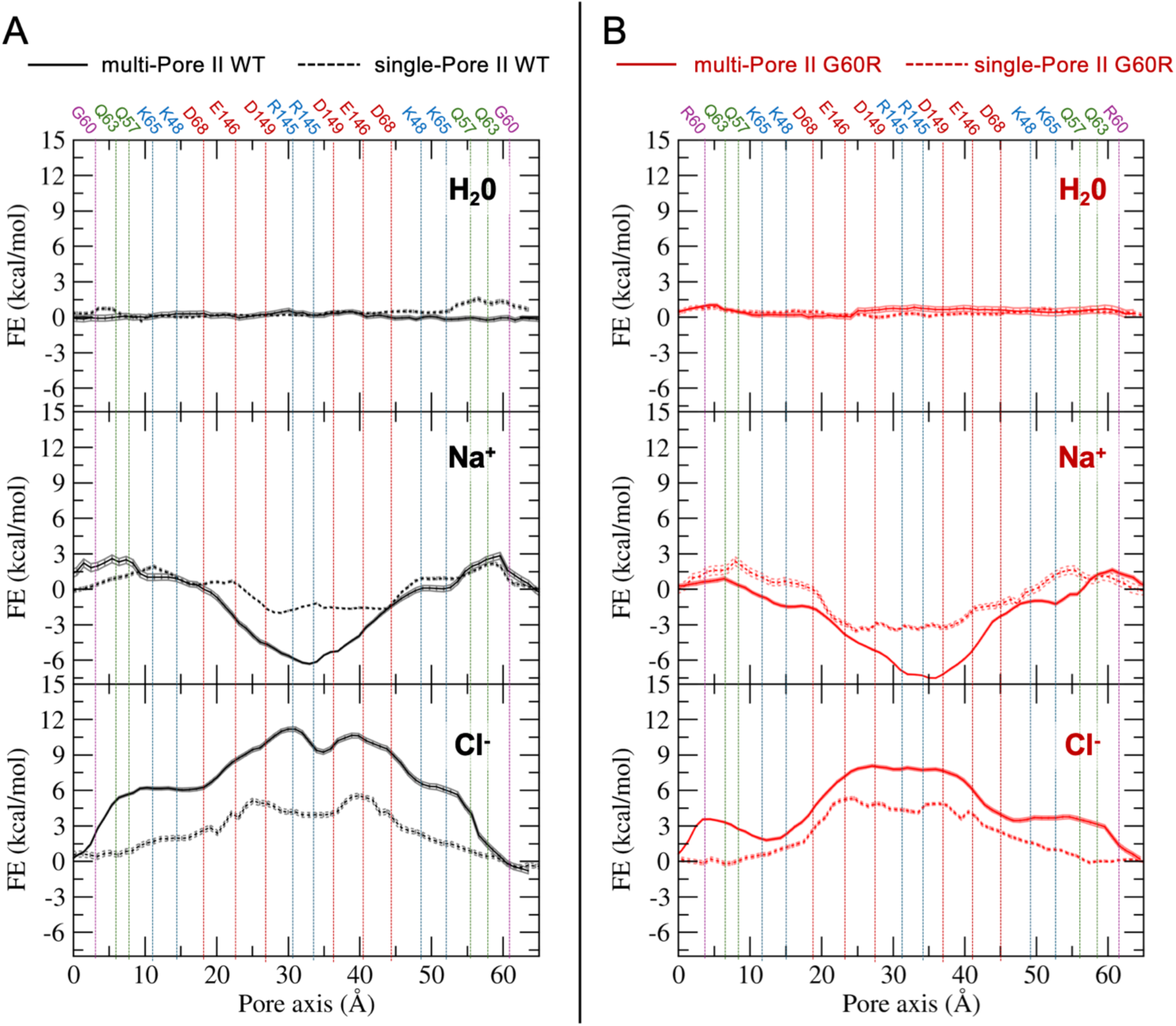
FE profiles for water and ion permeation across the BBB TJ models. **A**, WT multi-Pore II (solid black line), compared with single-Pore II^14^ (dashed black line). **B**, G60R multi-Pore II (solid red line), compared with single-Pore II^16^ (dashed red line). Vertical lines indicate the positions of the Cα atoms of the pore-forming residues.

Although water diffusion across BBB occurs at a relatively slow rate, to the best of our knowledge, there is no evidence to exclude the contribution of the paracellular pathway in water transport through the BBB. Consistently, previous computational investigations of Cldn5 strands always revealed water-permeable cavities^13,14,36–38^.

In the WT multi-Pore I, the Na^+^ ion profile showed an FE barrier of ∼ 7 kcal/mol in the central region of the pore (**Figure 8A**, middle plot). A similar result (barrier of ∼ 6 kcal/mol) was obtained in a control simulation using standard atomic masses and a time-step Δ*t* = 2 fs (**Figure S12**). Interestingly, the barrier was higher than the one obtained for single-Pore I. In the case of Cl*^−^*(**Figure 8A**, bottom plot), the system showed two barriers of ∼ 4.5 kcal/mol at each pore entrance (pore axis coordinate *y*= 10 Å and 50 Å), in correspondence with the negatively charged residues D68, E146, D149. In line with the presence of barriers for both cations and anions, standard MD simulations of the WT Cldn5 multi-Pore I system showed no evidence of charge transit through the paracellular cavities.

When considering the G60R mutant of Cldn5, the Na^+^ barrier was almost doubled (∼ 12 kcal/mol, **Figure 8B**, middle plot), as the mutation in each Cldn5 subunit generates a positively charged region in the middle of the paracellular cavities. Calculations for Cl*^−^*, on the other hand (**Figure 8B**, bottom plot), showed that the mutation results in an FE well about 7 kcal/mol deep in the middle of the cavity, canceling out the effect of the peripheral acidic residues and making the pore attractive for the anion, as demonstrated experimentally^15^. A similar result was obtained for the G60R single-Pore I, although the minimum was shallower, ∼3 kcal/mol deep. As for the multi-Pore II model, the WT system showed peripheral barriers and a central minimum for Na^+^ (**Figure 9A**, middle panel). Inserting the G60R mutation caused a reduction of the barriers and had almost no effect on the minimum depth, making the pore attractive for cations. When considering Cl*^−^*, the WT system showed a repulsive double peak of ∼ 11 kcal/mol (**Figure 9A**, bottom panel), which decreased to ∼7 kcal/mol in the mutant, still a significant barrier to cross at the physiological temperature. As seen for the Pore I systems, the multi-Pore II profiles show features that are in line with those of single-Pore II but enhanced. Hence, the multi-Pore II model does not reproduce the experimentally demonstrated effect of the G60R mutation. By looking at its structure (**Figure S11**), the substituted side chains are located at the periphery of the cavity and point outside of it, and their impact on the paracellular transport is marginal.

## Discussion

Assemblies of Cldn5 proteins in the BBB TJs are essential complexes to guarantee brain homeostasis. Because of that, it is not surprising that Cldn5 has become a promising target in the field of drug discovery. The development of engineered molecules able to transiently open BBB TJs and regulate the traffic to the brain may lead to unprecedented improvements in the treatment of various disorders. However, the absence of experimental data on Cldn-based quaternary structures limits the knowledge of these critical assemblies. Similarly to other applications, computational approaches can contribute with structural models that can be used, for example, to understand the impact of pathogenic mutations on the system’s structure and function. The recent report of the first Cldn5 variant linked to a neurological condition, G60R^15,39,40^, converting the TJ barrier into anion-selective channels via a modification in the ECL1 domain, provides an interesting test case for validating the models. In the last decade, several models of Cldn-based TJ assemblies were proposed^6^, two of which (called Pore I and Pore II) have been extensively studied for Cldn5^10,13,14,16^. We and other groups investigated these topologies as single, isolated paracellular pores formed by four protein subunits, an ideal system comprising the minimal functional unit and hence computationally convenient. However, this approach has limitations. First, the lack of adjacent protomers introduces potential inaccuracies due to an overestimated degree of hydration and larger flexibility of the paracellular loops that may affect transport properties. Second, in multi-pore structures the extracellular loops might adopt specific conformations that are not seen in single-pore ones. This is particularly relevant for the segment of ECL1 that is unresolved in both the crystal structure of Cldn15^35^ and the Suzuki model^12^, as it was investigated for Cldn10b in Refs.25 and 41. Finally, Cldn single-pore systems do not include identified protein-protein contacts such as the *cis-linear* pattern and some *trans-*interactions.

The role of extended polymerization was previously investigated for several members of the claudin family, namely Cldn3^42^, Cldn11^43^, Cldn15^20,22–24^, Cldn10a^26^ and Cldn10b^25,26,41,42,44^, but not for Cldn5. In this work, we studied two Cldn5 multimeric systems based on the Pore I (Suzuki, or JDR model) and Pore II topologies. By employing the HMR strategy to simulate with a time-step of 4 fs, we studied in all-atom details the dynamics of Cldn5 strands of sixteen protomers on the microsecond time scale and computed the FE profiles for single-ion permeation across the pores. Although TJs regulate the passage of multiple ions in the extracellular fluid, the single-ion FE profiles are a useful concept in permeation models, as they provide the location of binding sites and repulsive barriers within the protein cavities^45–47^. Our results corroborate the ability of the Suzuki-like model (the multi-Pore I) to recapitulate a series of experimental properties of the BBB TJs, with particular reference to the persistence of *cis-* and *trans-*protein-protein interactions, the ability to hinder the passage of ions, and the formation of aberrant anion channels once the G60R mutation is inserted. The FE barriers obtained for the multi-Pore I system are higher than those of the single-Pore I, revealing the importance of considering extended strands when studying paracellular transport.

A growing body of research supports the Suzuki architecture as the one capable of recapitulating the properties of both channel-forming and barrier-forming TJs. Recently, we showed that it can reproduce the differential permeability of Cldn10b for ions and water^41^, and employed it to design Cldn5-binding peptides^31^. In the future, further advancements can be made by applying deep-learning-based strategies to build the multimeric assemblies of claudins. In addition, the computational validation of the multi-Pore I model would certainly benefit from studies employing other force-fields different from CHARMM36m^48–51^ used in this and other works on paracellular channels^9,10,13,14,17–24^, and from the use of different membrane models^9,52^. In conclusion, our investigations identified a reliable structural model for Cldn5-based TJ strands, capable of reproducing BBB properties under physiological and pathological (the hemiplegia-associated Cldn5 G60R mutant^15^) conditions. In addition to advancing the knowledge of the BBB structure and function, the model can be used as a reference when designing novel Cldn-binding compounds for therapeutic applications^53–56^. Indeed, a detailed mapping of the protein-protein interfaces responsible for the stability of the assembly may help the identification of perturbing molecules or peptides to open the barrier and permit transient drug delivery^31^.

## Methods

### Homology modeling of Cldn5 proteins

Because of the lack of experimental structures, the human Cldn5 (UNIPROT: O00501) protein was built by homology modeling with SWISS-MODEL^57^, and using the crystal structure of the mouse Cldn15 protomer as a template (PDB ID: 4P79)^35^. Claudin proteins fold in a left-handed TM helix bundle that spans the membrane bilayer. The four TM *α*-helices are named TM1 to TM4. TM1 and TM2 are linked by a first long extracellular domain (ECL1), arranged into a four-stranded *β*-sheet (labeled as *β*1-*β*4), and stabilized by a disulfide bond conserved among various claudin proteins and formed by two cysteine residues in *β*3 and *β*4. In addition, a shorter extracellular helix (named ECH) is observed between *β*4 and TM2. Furthermore, TM3 and TM4 are connected by a second extracellular loop (ECL2), which is shorter than ECL1 (∼25 amino acids) and includes a further *β*5 sheet. Our homology-based Cldn5 structures, including all details of the homology modeling protocol, were presented in Refs. 14,16. A representative configuration of the Cldn5 protein with labeled domains is shown in **Figure S1**.

### Multi-Pore I models

To build the starting conformation of the WT Cldn5 multi-Pore I model, we used the coordinates of the multi-pore structure proposed by Suzuki et al. in Ref. 12 for the homologous Cldn15 (also named JDR model in Ref. 58 and in other works^25,26,42,44^). In this structure, the protein monomers interact via two distinct cis-interfaces, named linear and face-to-face59. The first one is the same observed in the crystal lattice of Cldn15^12^, while the second one is formed via interactions between the edges of *β*4-strands of ECL1 of side-by-side protomers. The *trans-*association between antiparallel claudin double rows from the membranes of adjacent cells results in a sequence of *β* barrel-like pores in the paracellular space. Our architecture comprises 16 Cldn5 subunits assembled to form three parallel pores.^12,58^ The previously described Cldn5 G60R mutation^15^ was introduced in each Cldn5 monomer using the rotamers tool available in UCSF Chimera^60^. Two versions of the model were considered, named G60R-A and G60R-B. In the former, the mutated side chains of the central pore monomers point away from the cavity, toward the lumens of the adjacent pores, while in the latter they point inside the inner pore. Before MD simulations, the quality of each model was improved using the GalaxyRefineComplex server^61,62^. We then relaxed the structures by minimizing their energy for 10 ps and equilibrating for 15 ns with a Generalized Born Implicit Solvent Simulation (GBIS)^63^ at 310 K, using NAMD/CHARMM36 and a time-step of 2 fs, and applying harmonic restraints on the atoms not belonging to the paracellular domains.

### Multi-Pore II models

The multi-Pore II architecture was built from the single-Pore II structure originally introduced in Refs. 9,10 and formed by *trans-*association of *cis-*dimers stabilized by a leucine zipper at the level of the TM2 and TM3 helices, supported by two homophilic *π*-*π* interactions between F127 and W138 on two opposing Cldn5 TM domains. This interaction was named back-to-back in Ref. 59. Since a template for a polymeric strand consisting of consecutive back-to-back dimers is currently unavailable, we generated the WT multi-Pore II model by replicating our previous single-Pore II system^14^ in the direction perpendicular to the pore axis. We first aligned two single-pores to form an octameric double-pore structure to obtain a repeated unit of minimal size but still containing surfaces between neighbor pores. From this, we selected two *cis-*dimers *trans-*interacting via their ECL2 domains diagonally across the two pores (i.e., one dimer from each pore) and used them as a repeated unit. We thus generated a three-pore strand by alternating manual repositioning and structural refinement with GalaxyRefineComplex^61,62^. Once the full 16-meric structure was obtained, it was relaxed by means of a preliminary simulation in an implicit solvent with the same protocol described for the multi-Pore I model. The resulting system comprises two antiparallel rows of Cldn5 interacting within the same membrane via a back-to-back *cis-*interface and a modified *cis-*linear interface. The G60R mutation was introduced in each Cldn5 monomer using the rotamers tool available in UCSF Chimera^60^.

### MD setup and parameters

Following the protocol of our previous works^14,16^, all the structural models of this work (multi-Pores I WT, G60R-A, G60R-B, and multi-Pores II WT, G60R) were further modified before MD simulations. Missing hydrogen atoms were added with CHARMM-GUI^64–66^, and disulfide bonds were assigned to the ECL1 cysteines, following the information provided by the experimental Cldn15 structure identified by the PDB ID: 4P79^35^. Then, each complex was inserted in a double membrane bilayer formed by 1-palmitoyl-2-oleoyl-SN-glycero-3-phosphocholine (POPC) molecules to reproduce the native state of two facing cells in TJs, using VMD psfgen^67^, and solvated with explicit water molecules and ions.

All simulations were performed with the NAMD software^68^ in the NPT ensemble (P=1 atm, T=310 K) by employing the Nosé-Hoover Langevin piston method^69,70^ and a Langevin thermostat. The hydrogen mass repartitioning (HMR) method was used to speed up calculations by allowing a time-step Δ*t* = 4 fs^32,33^. The oscillation period of the piston was set at 300 fs, and the damping time scale at 150 fs^34^. The Langevin thermostat was employed with a damping coefficient of 1 ps*^−^*^1^. The parameters provided by CHARMM36m^48–50–CHARMM3671^ were used for the protein and lipids, respectively. The TIP3P model was employed for water molecules^72^. The CHARMM ionic parameters with NBFIX corrections^73–75^ were used. Electrostatic and Van der Waals (VdW) interactions were calculated with the standard CHARMM cutoff of 12 Å. A switching function was applied starting to take effect at 10 Å to obtain a smooth decay. The VdW Force Switching option for CHARMM36 was applied^76^. Hexagonal periodic boundary conditions were used^14,16^. Long-range electrostatic interactions were calculated using the Particle Mesh Ewald (PME) algorithm^77^, by adopting a spline interpolation order 6. A maximum space between grid points of 1.0 Å was used. Covalent bonds including hydrogen atoms were constrained using the SHAKE/SETTLE algorithms^78,79^. Electrostatic and VdW interactions were computed at each simulation step^14,34^. In the simulations of the multi-Pore I model, we applied positional restraints on the following atoms:

- The C*α* atoms at the termini of TM helices in each Cldn5 monomer, i.e. of residues E7, G10, L13, G17, G20, L23, Q78, R81, V85, V93, F96, L99, G111, K114, V117, W138, N141, G161, L164, G167, L174, G177, L181.
- The C*α* atoms of the ECL domains of the peripheral protomers (at the end of the strands), i.e., residues P28 to L73 and D149 to G161.

Restraints are hence applied to selected atoms of the TM helices and of the paracellular domains at the periphery of the structures, as we did in previous works^14,16,17,19^. Their use is justified by considering that, despite including multiple monomers, our TJ models still lack the extended strand framework and cytoskeletal tethering via ZO proteins, which limit their spatial fluctuations in the physiological environment.^24,80^. In **Figure S2,** we show images of the multi-Pore I and multi-Pore II, respectively, with the restrained C*α* atoms represented as red spheres.

### Control simulations

We performed two control simulations of the multi-Pore I (WT) using standard atomic masses and time-step Δ*t* = 2 fs. In the first one, the same set of restraints described above was applied, hereafter referred to as *extended*, while in the second one, only the C*α* of the most peripheral Cldn5 monomers were fixed (*restricted* set). The system was prepared as the HMR ones, and the NPT ensemble (P = 1 atm and T = 310 K) was ensured in NAMD^68^ by applying the Nosé-Hoover Langevin piston method^69,70^ with an oscillation period of 50 fs and a damping time scale of 25 fs, following the standard NAMD scripts available in CHARMM-GUI^64,65^. The Langevin thermostat was employed with a damping coefficient of 1 ps*^−^*^1^. In **Figure S2 A,** we show an image of the multi-Pore I with the restrained C*α* atoms of the restricted set represented as green spheres.

### Structural analysis

The MD trajectories were visualized and analyzed using VMD^67^ (www.ks.uiuc.edu/Research/vmd/) with Tcl scripts and UCSF Chimera^60^ (www.cgl.ucsf.edu/chimera/). The NAMD-COLVAR module^81^ was also used for analysis.

#### Pore radius calculation

We used the HOLE program^82,83^ to calculate the pore radius of each model. Snapshots were extracted from simulated trajectories every 25 ns, and the resulting protein structures were used to calculate the radius. The frames belonging to the first 250 ns were excluded in each simulation. Data for each of the three pores were lumped together to obtain a single average profile with standard deviation. Pictures were produced using VMD.

#### RMSD calculations

Different sets of atoms were used to calculate the backbone Root Mean Squared Deviations (RMSDs) from initial structures (**Figure S3**). The RMSD of the full system (ECLs + TM domains) was calculated considering all protomers except for the peripheral ones. The RMSDs of individual pores were calculated using the backbones of the ECLs only (residues P28 to V77 and D149 to G161 of all protomers).

### Calculations of *cis-* and *trans-*interactions distances

To test the stability of the multi-pore assemblies, we monitored a set of distances between protein atoms. We started with residues involved in the *cis-*linear interactions, first observed in the crystallographic structure of the Cldn15 homolog (PDB ID: 4P79)^35^ and essential in maintaining the row arrangement of the strand^12^. We thus considered four distances (d1 to d4) between Cldn5 residues corresponding to those defining the *cis-*linear interface in Cldn15. Four additional distances (d5 to d8) were monitored between monomers in *trans-*interactions within the multi-Pore I model, with the last one being measured only in the standard atomic masses control simulations.

#### Stability of the *β* barrel-like paracellular cavities in the multi-Pore I

A set of additional distances was monitored to assess the integrity of the paracellular *β*-barrel in the multi-Pore I. Specifically, we used the pairs of V154 and V70 found at the entrances of each cavity and the couples of Q57 and of Q63 pairs at the center of the paracellular space. All distances were calculated using the C*α* atoms or the most external carbon atom of the side chains (the C*β* and C*δ* atoms of the valine and glutamine residues, respectively).

### Umbrella sampling simulations with extended set of TM restraints

We used the umbrella sampling (US) method to calculate one-dimensional (1D) FE profiles for the permeation of the main physiological ions (Na^+^ and Cl*^−^*) and one H_2_O molecule through the central paracellular pores. We applied the same protocol as in our previous works^14,16,18,19^. Briefly, a harmonic potential energy term is added to the MD force field to enhance the sampling along the selected collective variable (CV) in different and independent simulations (named *windows*). Here, we used the projection of the ion (or water molecule) position onto the axis of the paracellular channel, which is oriented along the *y*-axis. The potential added in window *i* is expressed as 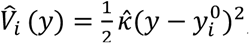, where 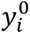 indicates the value at which the CV is restrained, and *κ̂* is a constant. We performed US simulations with these harmonic restraints and *κ̂* = 2 kcal/(mol*·*Å^2^), allowing sufficient overlap between the sampled distributions of adjacent windows. We employed a total of 70 windows with a uniform spacing of 1 Å, mapping from *−*35 to +34 Å along the *y-*axis. In each window, after the minimization, a preliminary equilibration of 2 ns was performed (1 ns using a 2-fs time-step and the second one with 4 fs). Several starting conformations were produced by swapping a few equilibrated water molecules with the tested ion inside the window. The time required for convergence for each window is shown in **Table 2**. All ions but the one used in the CV were excluded from the paracellular cavity of the central pore using two half-harmonic potentials, one for each entrance, with an elastic constant of 10 kcal/(mol*·*Å^2^). The displacement of the ion orthogonal to the pore axis was confined within a disk of radius *r*_0_ + *δ*, where *r*_0_ is the pore radius as determined by the HOLE program^82,83^ and *δ* = 2 Å, by means of a potential energy term and a force constant of 2 kcal/(mol*·*Å^2^)^14^. Simulations were performed with the HMR method and time-step Δ*t* = 4 fs in the NPT ensemble with the same algorithms and parameters discussed above. A control US calculation was performed for the permeation of Na^+^ through the multi-Pore I using standard atomic masses and time-step Δ*t* = 2 fs, in the NPT ensemble with the same parameters as the standard masses simulation in the previous section.

**Table 2.**
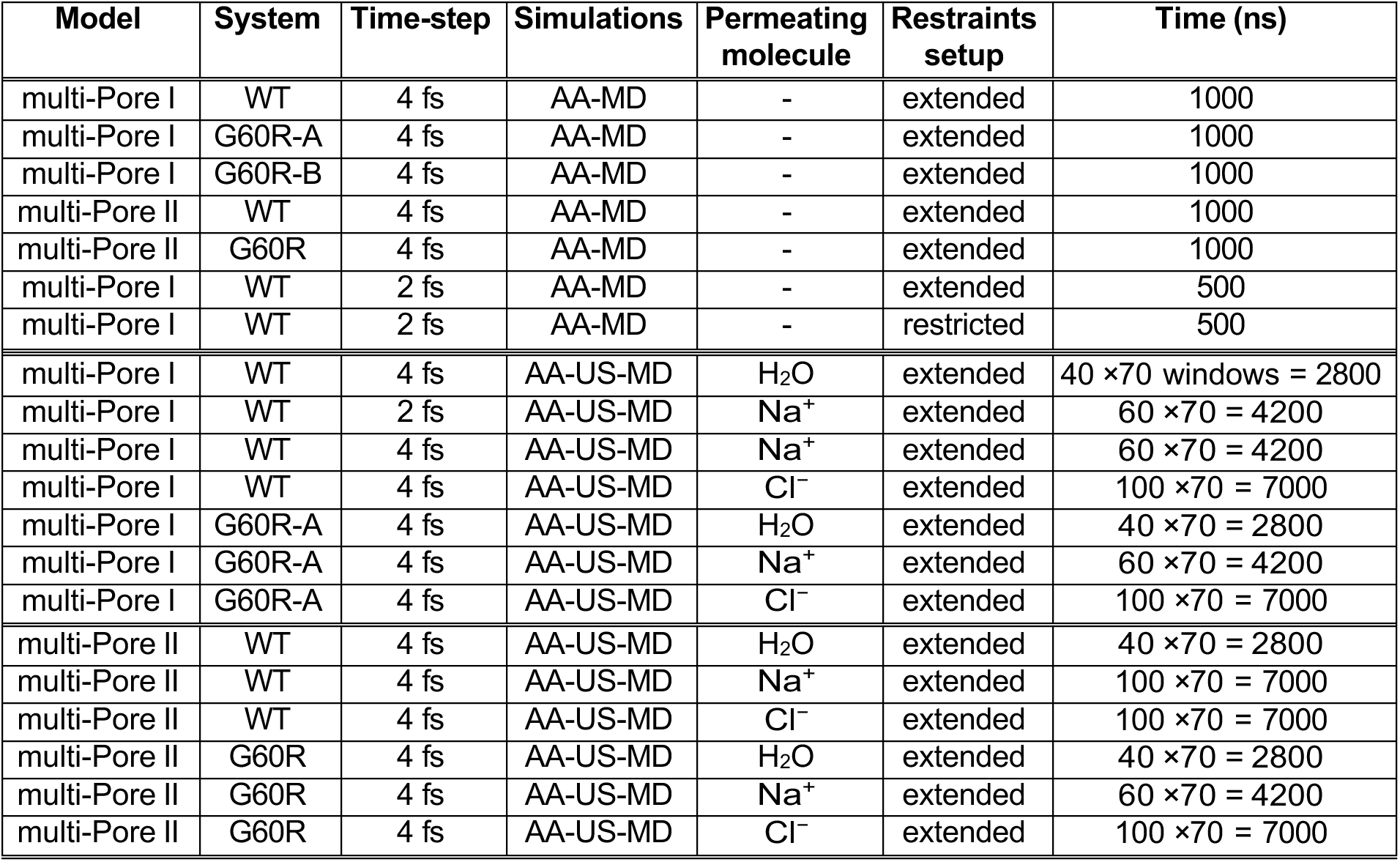
Summary of the MD simulations performed in this work.

| Model | System | Time-step | Simulations | Permeating molecule | Restrains setup | Time (ns) |
| --- | --- | --- | --- | --- | --- | --- |
| multi-Pore I | WT | 4 fs | AA-MD | - | extended | 1000 |
| multi-Pore I | G60R-A | 4 fs | AA-MD | - | extended | 1000 |
| multi-Pore I | G60R-B | 4 fs | AA-MD | - | extended | 1000 |
| multi-Pore II | WT | 4 fs | AA-MD | - | extended | 1000 |
| multi-Pore II | G60R | 4 fs | AA-MD | - | extended | 1000 |
| multi-Pore I | WT | 2 fs | AA-MD | - | extended | 500 |
| multi-Pore I | WT | 2 fs | AA-MD | - | restricted | 500 |
| multi-Pore I | WT | 4 fs | AA-US-MD | H<sub>2</sub>O | extended | 40 × 70 windows = 2800 |
| multi-Pore I | WT | 2 fs | AA-US-MD | Na<sup>+</sup> | extended | 60 × 70 = 4200 |
| multi-Pore I | WT | 4 fs | AA-US-MD | Na<sup>+</sup> | extended | 60 × 70 = 4200 |
| multi-Pore I | WT | 4 fs | AA-US-MD | Cl<sup>-</sup> | extended | 100 × 70 = 7000 |
| multi-Pore I | G60R-A | 4 fs | AA-US-MD | H<sub>2</sub>O | extended | 40 × 70 = 2800 |
| multi-Pore I | G60R-A | 4 fs | AA-US-MD | Na<sup>+</sup> | extended | 60 × 70 = 4200 |
| multi-Pore I | G60R-A | 4 fs | AA-US-MD | Cl<sup>-</sup> | extended | 100 × 70 = 7000 |
| multi-Pore II | WT | 4 fs | AA-US-MD | H<sub>2</sub>O | extended | 40 × 70 = 2800 |
| multi-Pore II | WT | 4 fs | AA-US-MD | Na<sup>+</sup> | extended | 100 × 70 = 7000 |
| multi-Pore II | WT | 4 fs | AA-US-MD | Cl<sup>-</sup> | extended | 100 × 70 = 7000 |
| multi-Pore II | G60R | 4 fs | AA-US-MD | H<sub>2</sub>O | extended | 40 × 70 = 2800 |
| multi-Pore II | G60R | 4 fs | AA-US-MD | Na<sup>+</sup> | extended | 60 × 70 = 4200 |
| multi-Pore II | G60R | 4 fs | AA-US-MD | Cl<sup>-</sup> | extended | 100 × 70 = 7000 |

In all US simulations, we employed the extended setup of restraints described above, thus avoiding orientational and translational displacements of the proteins that could affect FE calculations. Residues in the paracellular space were not restrained. The Weighted Histogram Analysis Method (WHAM)^84,85^ was used to reconstruct the 1D-FE, using the code implementation provided by the Grossfield group, available at http://membrane.urmc.rochester.edu/content/wham. For this analysis, we selected 70 bins and a tolerance of 0.00001. Statistical uncertainty was evaluated in each bin via bootstrapping with 100 trials^18^.

## Main abbreviations and notations

AA: All Atom
BBB: Blood-Brain Barrier
CLDN: Claudin
COM: Center Of Mass
CV: Collective Variable
ECL: ExtraCellular Loop
FF: Force Field
FE: Free Energy
GBIS: Generalized Born Implicit Solvent
HMR: Hydrogen Mass Repartitioning
JDR: Joined anti-parallel Double-Row model
MD: Molecular Dynamics
PME: Particle Mesh Ewald
POPC: 1-Palmitoyl-2-Oleoyl-sn-glycero-3-PhosphoCholine
TJ: Tight Junction
TM: TransMembrane
US: Umbrella Sampling
VdW: Van der Waals
WHAM: Weighted Histogram Analysis Method
WT: Wild Type

## Author Contributions Statement

- **Conceptualization:** Giulio Alberini and Luca Maragliano.
- **Data curation:** Alessandro Berselli, Giulio Alberini, Linda Cerioni.
- **Formal analysis:** Alessandro Berselli, Giulio Alberini, Linda Cerioni, Luca Maragliano.
- **Funding acquisition and resources:** Luca Maragliano, Fabio Benfenati.
- **Investigation:** Alessandro Berselli, Giulio Alberini, Linda Cerioni, Luca Maragliano.
- **Project administration:** Alessandro Berselli, Giulio Alberini, Luca Maragliano, Fabio Benfenati.
- **Supervision:** Alessandro Berselli, Giulio Alberini, Luca Maragliano.
- **Visualization:** Alessandro Berselli.
- **Writing - original draft:** Giulio Alberini.
- **Writing - review & editing:** Giulio Alberini, Alessandro Berselli, Linda Cerioni, Luca Maragliano and Fabio Benfenati.

## Acknowledgments

We gratefully acknowledge the Data Science and Computation Facility and its Support Team for their support and assistance on the IIT High-Performance Computing Infrastructure. We thank Alessia Vignolo, Sergio Decherchi and Mattia Pini for their kind assistance. We acknowledge ISCRA for awarding this project access to the LEONARDO supercomputer, owned by the EuroHPC Joint Undertaking, hosted by CINECA (Italy). We also thank Diego Moruzzo, Ilaria Dallorto, Rossana Ciancio, and Arta Mehilli for administrative assistance and technical help.

## Competing Interests and ethics statements

The authors have no financial or non-financial interests to disclose. In this work, no potentially identifiable human images or data are presented

