## Supplementary Information for "A multi-pore model of the blood-brain barrier tight junction strands recapitulates the permeability features of wild-type and mutant claudin-5"

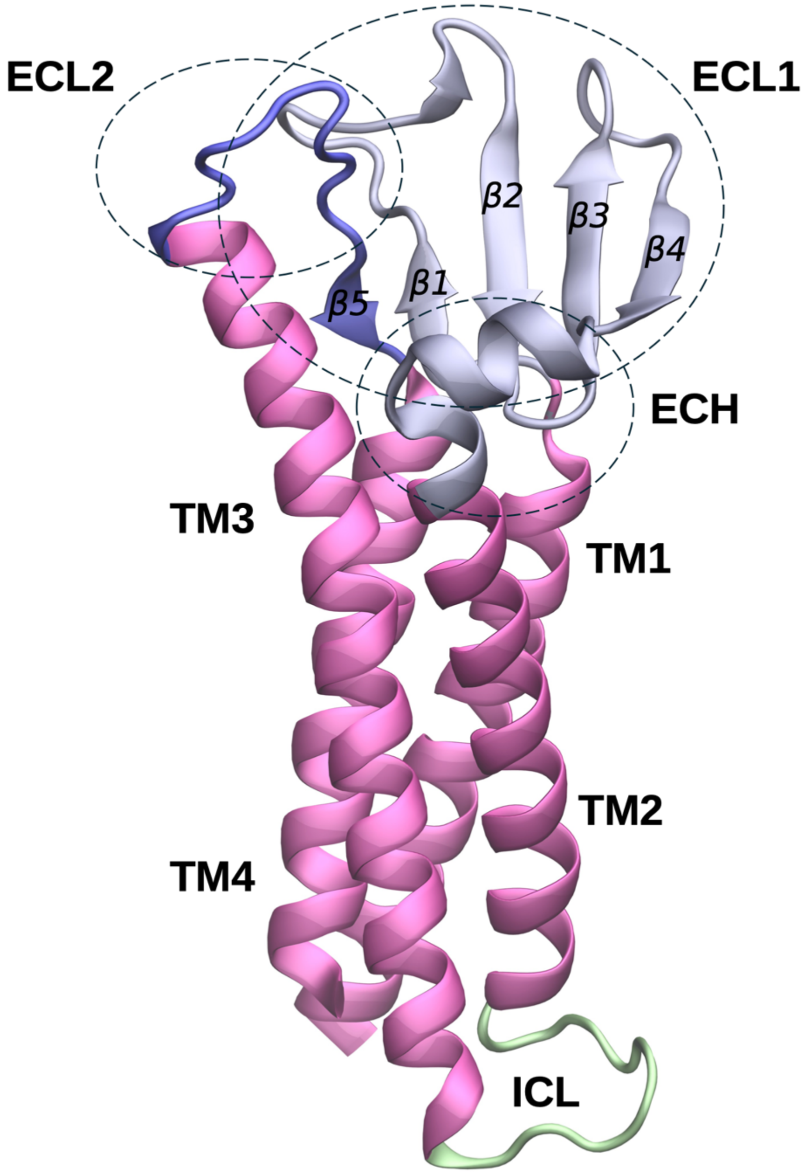


***Figure S1. Three-dimensional structure of the Claudin-5 monomer.*** *The three-dimensional structure of the Cldn5 monomer was reproduced with SWISS-MODEL (https://swissmodel.expasy.org) using the Cldn15 crystal as a template (PDB ID: 4P79). The structural domains are indicated with the nomenclature used in the main text and distinguished by their coloring: magenta for TM domains, silver for ECL1, blue for ECL2, and green for the intracellular loop (ICL).*

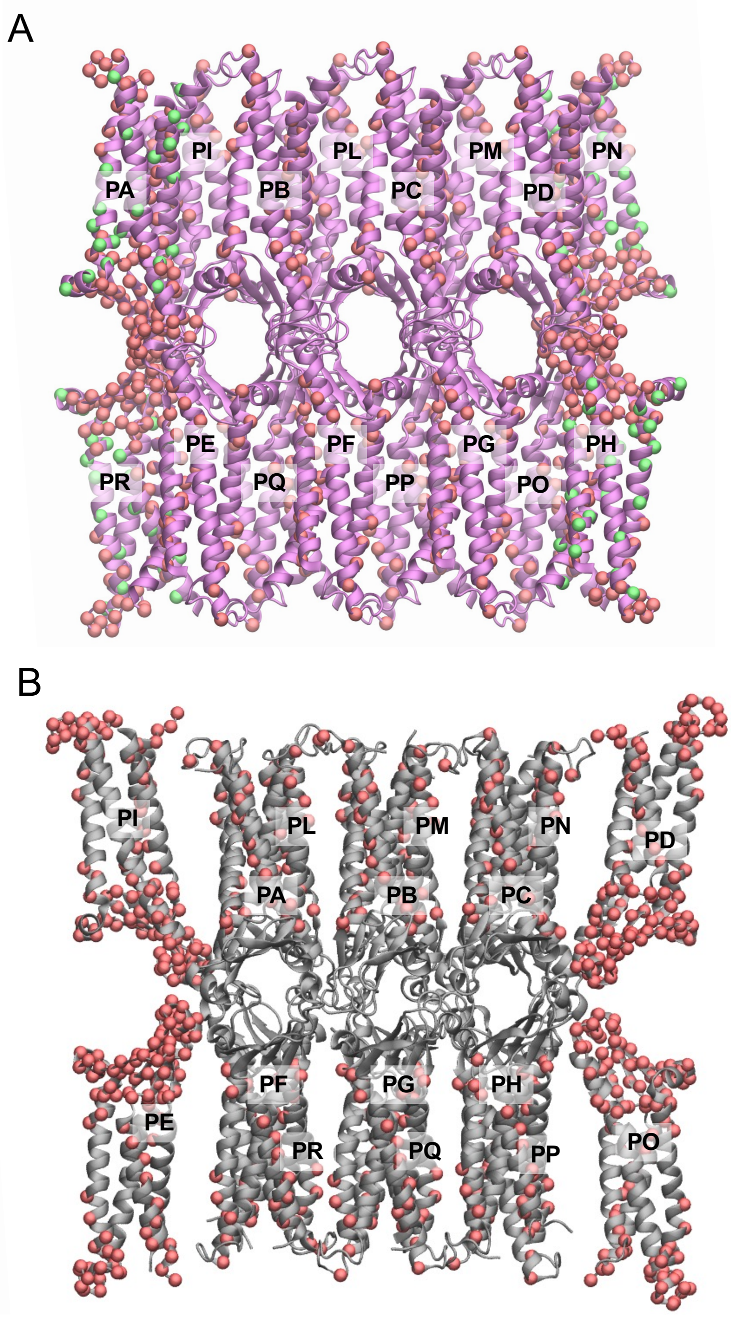


***Figure S2. Positional restraints used in MD simulations of multi-Pore models.*** *Positionally restrained Cα atoms are shown as VdW spheres colored in red for the extended set and green for the restricted set. (A) multi-Pore I. (B) multi-Pore II.*

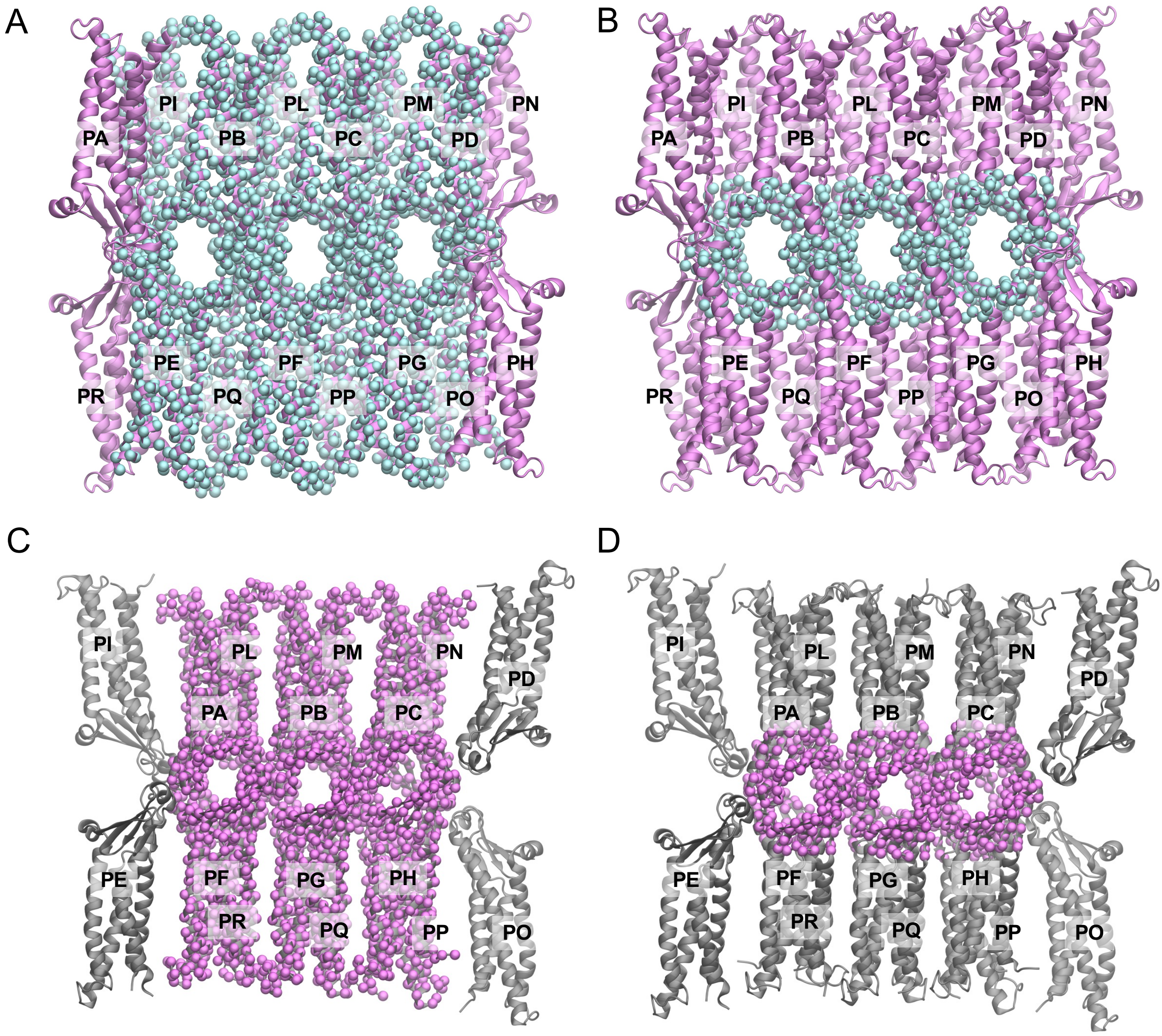


***Figure S3. Atoms considered in the RMSD calculations.*** *VdW spheres indicate the backbone atoms of the multi-Pore I (**A,** whole system, excluding peripheral protomers; **B,** paracellular domain) and multi-Pore II (**C,** whole system, excluding peripheral protomers; **D,** paracellular domain) models used for RMSD calculations.*

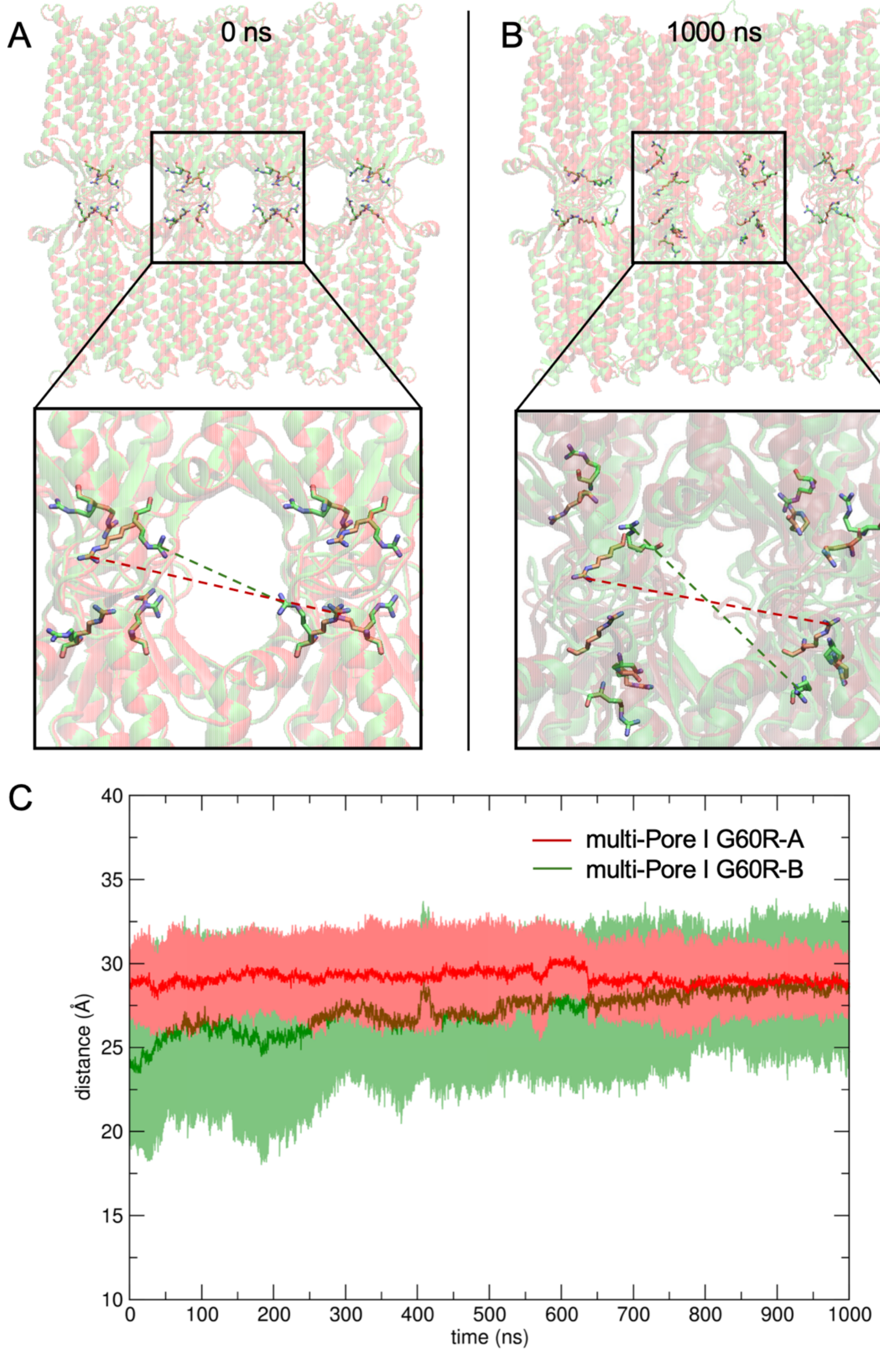


***Figure S4. Structural analysis of the G60R multi-Pore I models. A****, structures of the G60R-A and G60R-B multi-Pore-I like structures before MD simulations. For each panel, the R60 sidechains are shown in sticks.* ***B****, the same structures of the G60R-A and G60R-B systems at the end of the standard AA-MD simulations.* ***C****, average cross-distance between the R60 CZ-atom of multi-Pore I G60R-A (red trace) and G60R-B (green trace). The average value and associated error are calculated as the mean and standard deviation using each pair of facing R60 residues across the three pores of each system.*

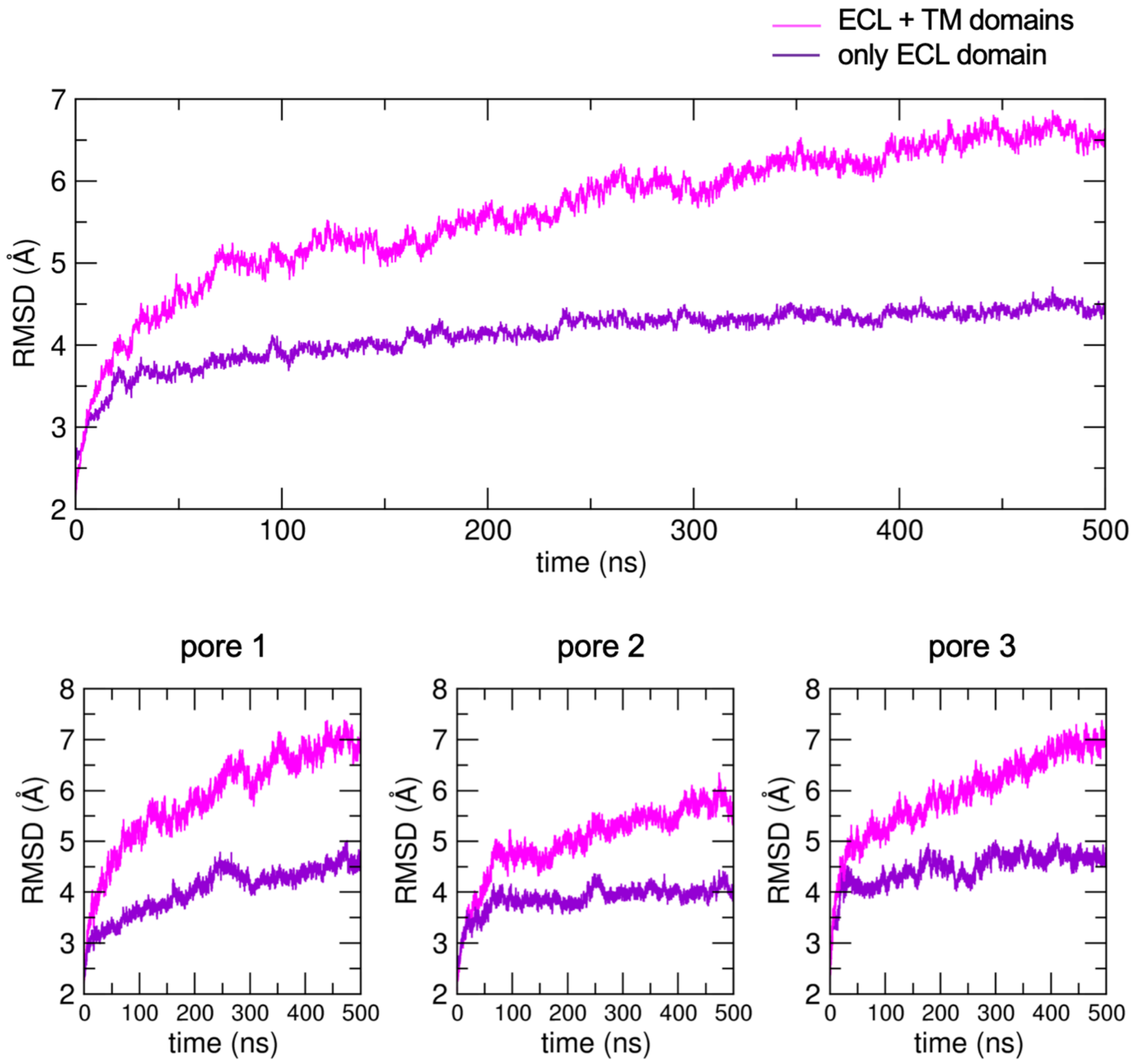


***Figure S5. RMSD of the inner protomers of the multi-Pore I WT, restricted set of restraints.*** *The RMSD of the protein backbone is calculated for the entire system in the upper panel and for the three individual pores included in the system in the lower ones. The RMSD is calculated including both the TM and the ECL domains (magenta profile) or considering only the ECL domain (purple profile). The four outermost protomers are kept fixed and excluded from the calculation of the RMSD, as described in the main text.*

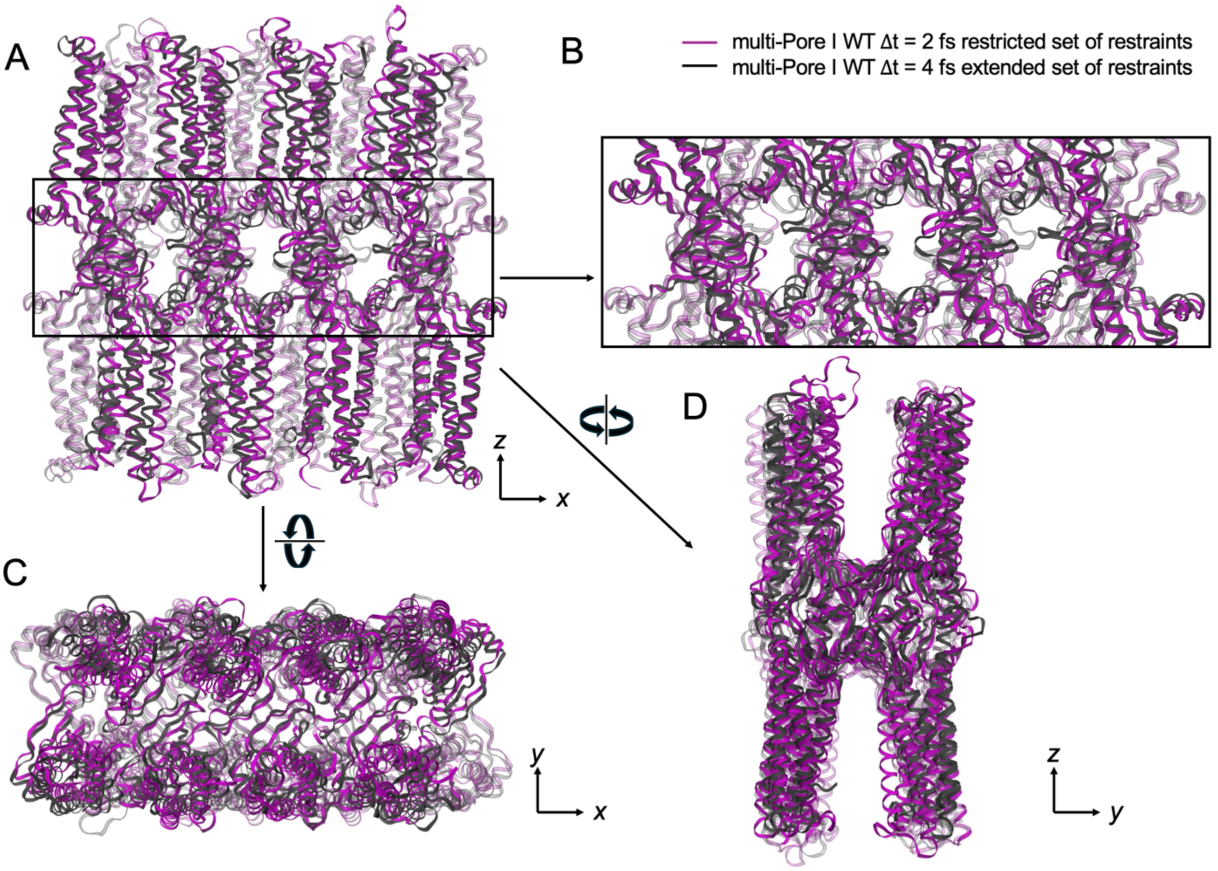


***Figure S6. Superposition of the Cldn5 multi-Pore I WT configurations simulated with and without positional restraints.*** *The configuration obtained after 500 ns of standard MD simulations with HMR (black) end extended set of restraints is superimposed to the configuration obtained after the same time interval in the simulations with standard masses and restricted set of restraints (purple). The systems are shown from the basolateral (**A, B**) and apical (**C, D**) views.*

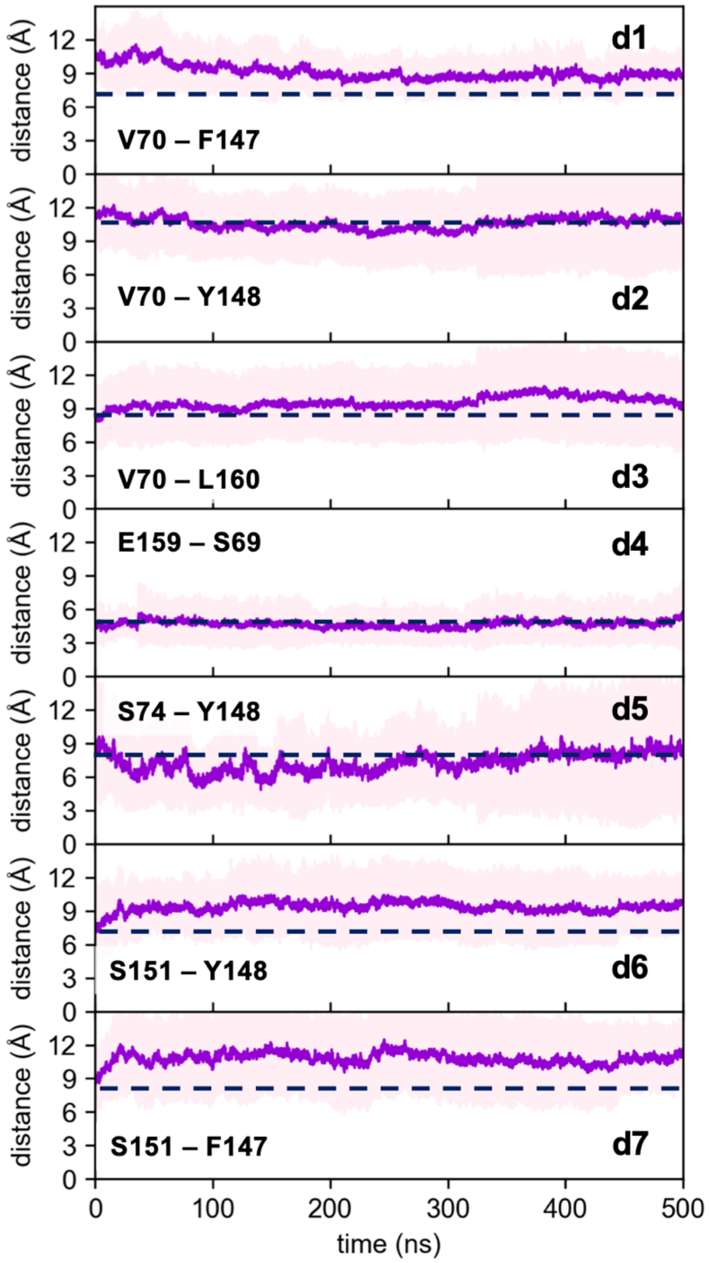


***Figure S7. Cis- and trans-interactions calculated for the Cldn5 multi-Pore I WT system simulated with the restricted set of restraints.*** *Distances are calculated on a timescale of 500 ns of standard MD simulations and named with the same nomenclature used in the main text. The average value of the distances calculated from MD simulations of Multi-Pore I WT with the extended set of restraints and Δt = 2 fs is reported as the blue dotted line.*

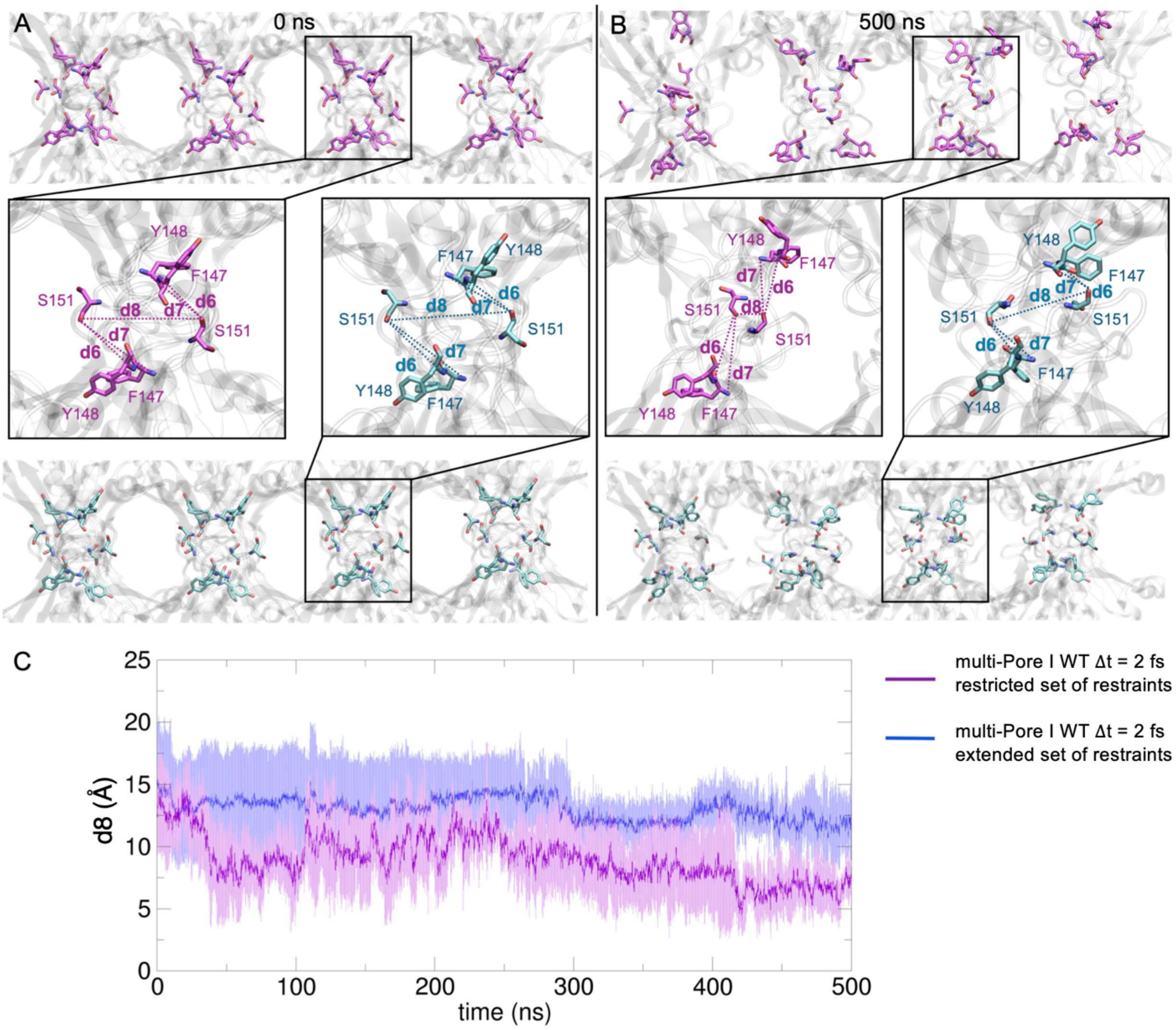


***Figure S8. Modification of the trans-interactions in the multi-Pore I models.*** *Distances between S151 and Y148 (d6), S151 and F147 (d7), and between opposing S151 residues (d8) are shown for the multi-Pore I simulated with the extended (blue structures) or restricted (purple structure) set of positional restraints and a time step of 2 fs. The configurations at the beginning and the end of the MD simulations are shown in panels **A** and **B**, respectively. **C**, evolution of the distance between the hydroxyl O-atoms belonging to the sidechains of opposing S151 residues (d8) in the restrained (blue profile) and unrestrained (purple profile) MD simulations. The profile and the associated error are calculated as average and standard deviation over each pair of S151 residues in the multi-Pore I models.*

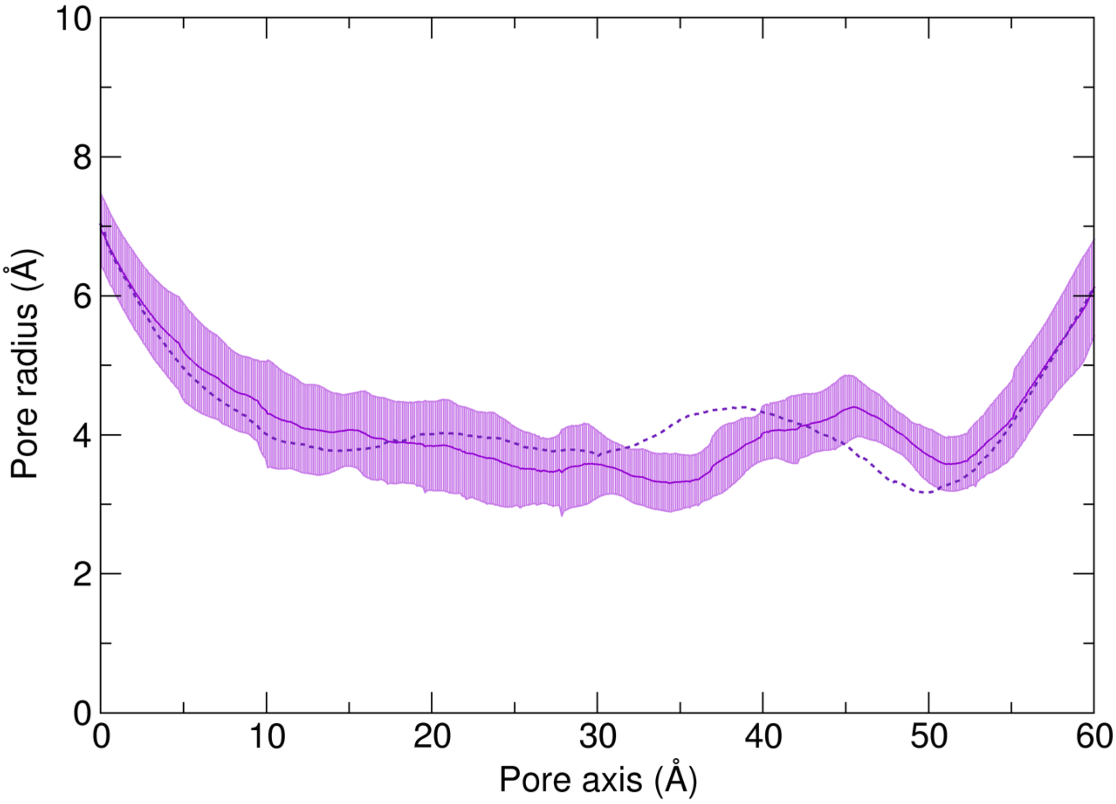


***Figure S9. Pore radius profiles of the Cldn5 multi-Pore I WT system simulated with the limited set of restraints.*** *Time-average of the central pore radius (solid purple line) and associated error (shaded area) were calculated using the HOLE program. The dotted purple line is the average profile calculated over the three pores of the system.*

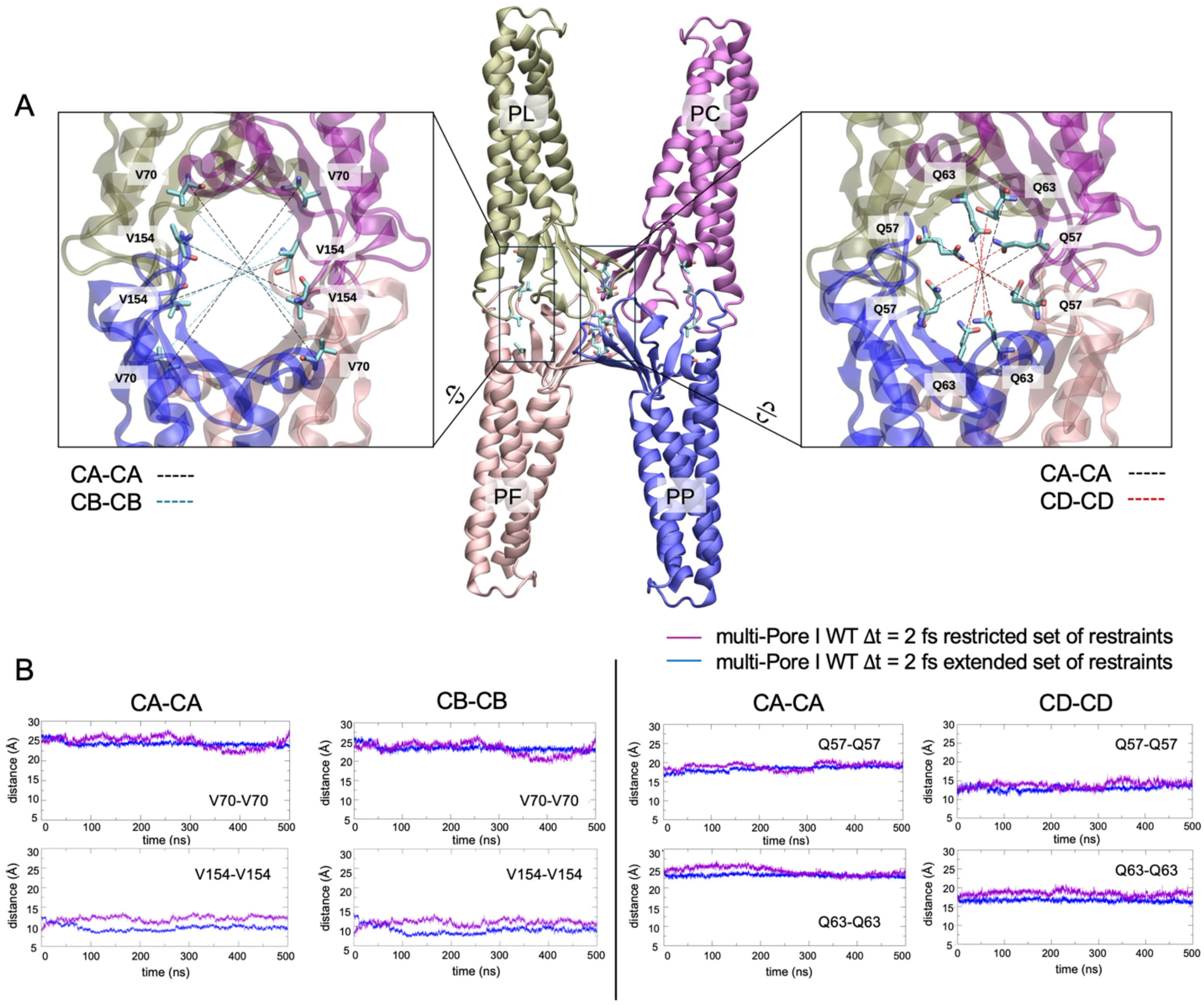


***Figure S10. Stability of the multi-Pore I β-barrel during the MD simulations. A****, three-dimensional representation of the central pore in the multi-Pore I WT model. Close-up views of the two pairs of residues considered to assess the integrity of the β-barrel are shown on the left for the V154 and V70 found at the entrances of the pore, and on the right for the Q57 and Q63 pairs at the center of the cavity.* ***B****, average distances were calculated during the MD simulations in the presence (blue) and absence (purple) of positional restraints and a time step of 2 fs. Distances are calculated using the Cα-atoms (CA) or the most external C-atom belonging to the sidechains (CB is the Cβ , and CD is the Cδ atom of the valine and glutamine residues, respectively).*

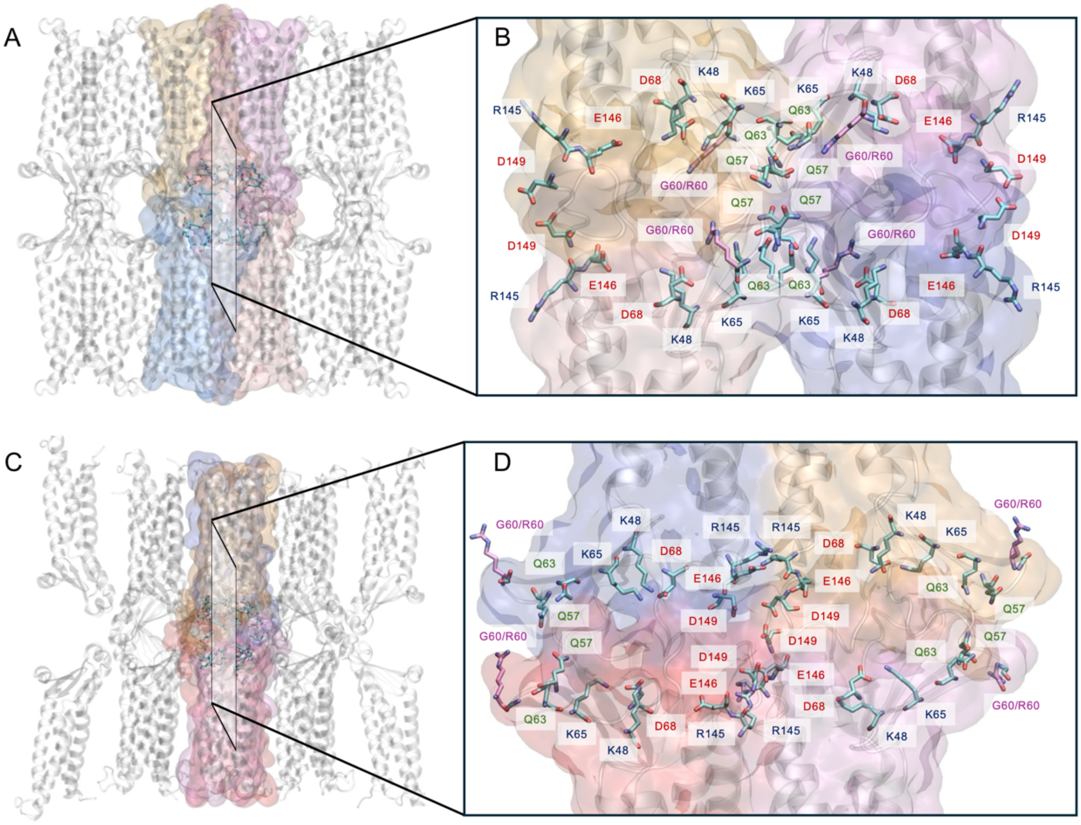


**Figure S11.** ***Pore-lining residues in multi-Pore I and multi-Pore II models. A***, *equilibrated structure of multi-Pore I. All the Cldn5 proteins are represented using gray ribbons. Colored surfaces are added to the four Cldn5 proteins forming the central pore.* ***B***, *a cross-section of the central paracellular cavity, with the side chains of the pore lining residues represented as sticks and the mutated G60R side chains colored in purple.* ***C***, *equilibrated structure of multi-Pore II. All Cldn5 proteins are represented using grey ribbons. Colored surfaces are added to the four Cldn5 proteins, forming the central pore.* ***D***, *cross-section of the central paracellular cavity, with the same representation of panel B.*

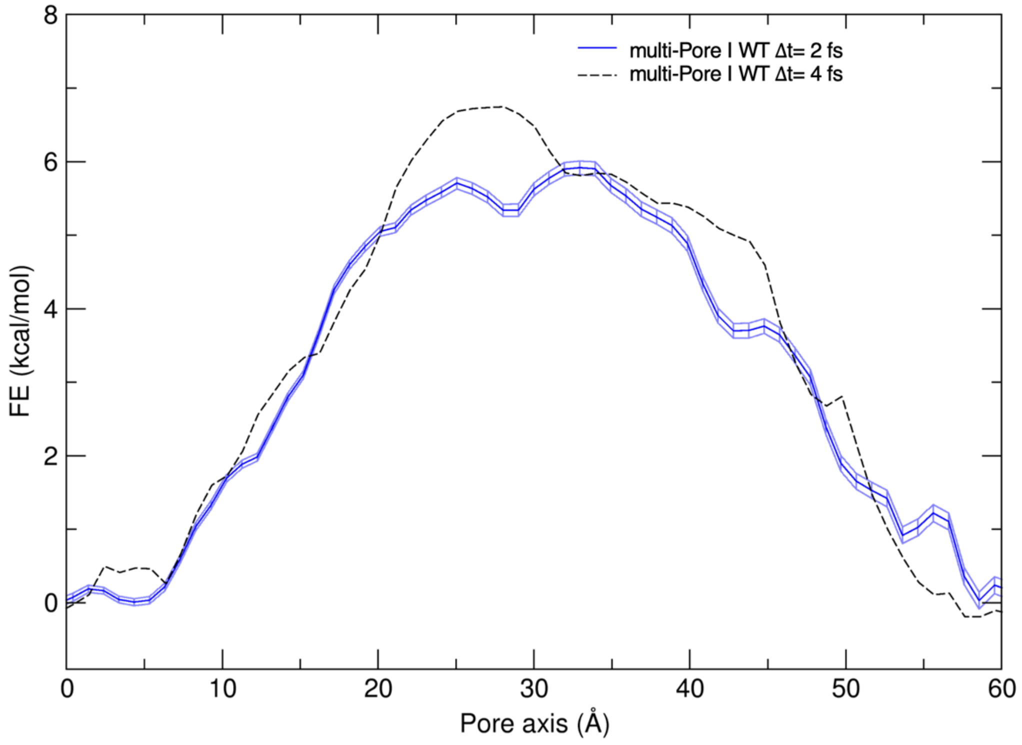


***Figure S12. Free energy profile of $Na^+$ permeation through the Cldn5 multi-Pore I performed with a time step of 2 fs.*** *A representative configuration was selected after 200 ns of standard MD simulation and used to perform the FE calculation with the US-WHAM method, adopting a time step of 2 fs and using the same protocol described for the multi-Pore I WT model discussed in the main text. The FE profile obtained with HMR is also provided (black dashed line) for comparison.*
